# A different world: temporal changes in nudibranch community structure over a half-century

**DOI:** 10.1101/2025.07.31.668013

**Authors:** Riko Kato, Mitsuharu Yagi

**Affiliations:** Graduate School of Fisheries and Environmental Sciences, Nagasaki University, Nagasaki 852-8521, Japan

**Keywords:** biological diversity, nudibranch, opisthobranchia, climate change

## Abstract

Understanding long-term changes in marine biodiversity is essential for evaluating effects of climate change on coastal ecosystems. In this study, we compared nudibranch assemblages in northwestern Kyushu, Japan, during three time periods (1960–1980, 2001–2003, and 2023–2024), based on underwater surveys and historical records. In all, 47 nudibranch species were recorded during 27 diving surveys conducted in 2023 and 2024. Species diversity indices (Shannon–Wiener H and Simpson D) showed higher values during this survey than in 2001–2003. Comparative analysis of species composition revealed significant shifts, with 15 species exhibiting statistically significant differences in relative abundance from the past to the present. Notably, several species common in the past, such as *Aplysia kurodai*, were rarely observed in the recent survey, while many tropical-subtropical species appeared for the first time. The proportion of tropical-subtropical species increased markedly, whereas subarctic species were no longer detected. Similarity indices (Jaccard’s coefficient and Sorensen–Dice index) indicated that the current community differs markedly from those in earlier periods. These findings suggest a major community reorganization, potentially driven by rising sea water temperatures and other environmental changes. This study highlights the importance of long-term monitoring using multiple indicators to detect and interpret climate-driven biodiversity shifts in coastal marine ecosystems.

**Highlights:**

- Long-term changes in nudibranch assemblages were investigated using visual census surveys spanning 6 decades.
- Species richness and tropical species have increased markedly in recent decades.
- Subtropical species expanded their ranges, while some cold-water species declined.
- Findings suggest that community-level tropicalization is likely driven by ocean warming.

**Graphical abstract:** 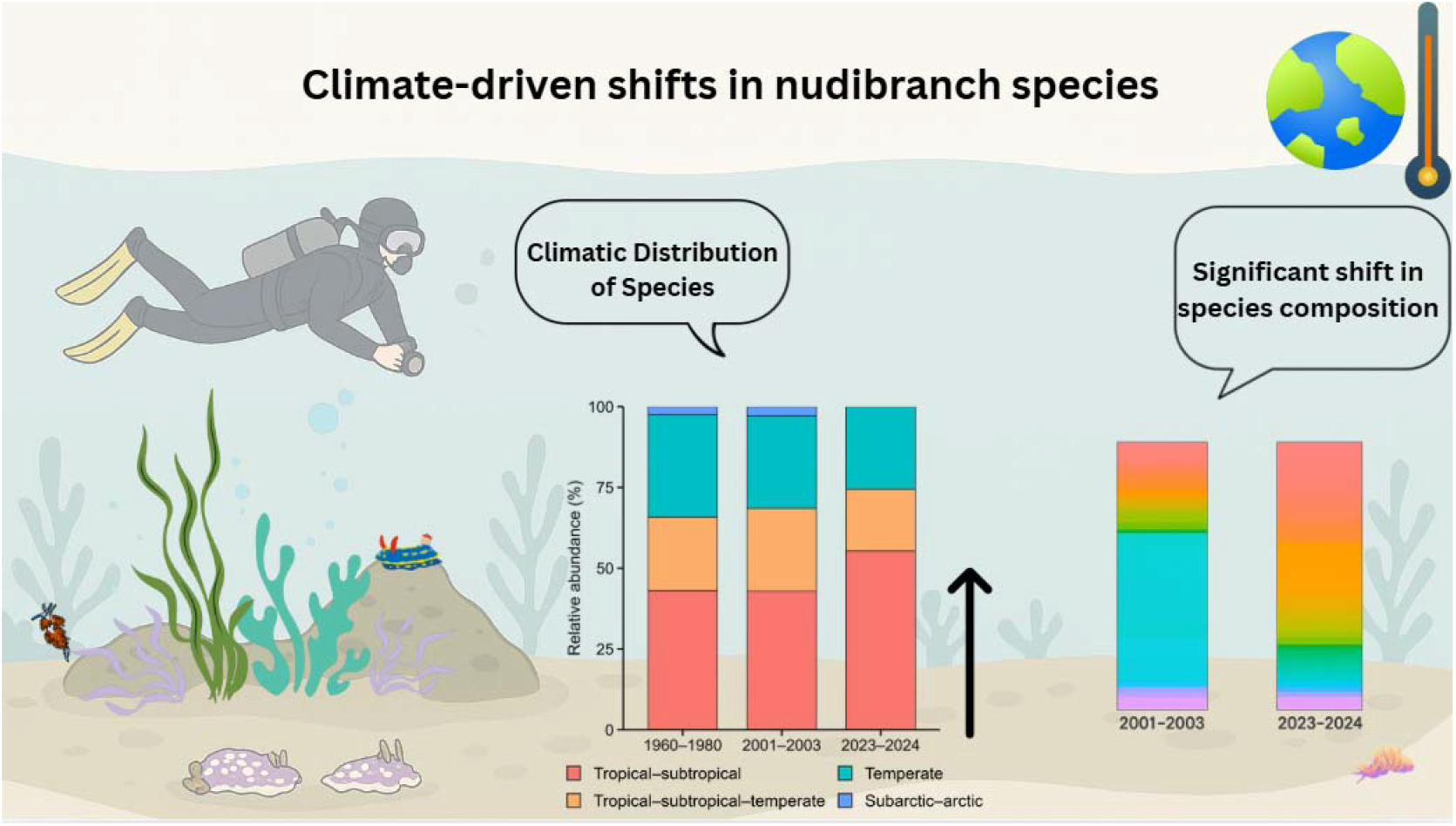

## 1. Introduction

Climate change has extensively impacted marine ecosystems, influencing ecosystem functions through shifts in biodiversity. Numerous previous studies have highlighted the serious effects of global warming and climate change on biological communities (Pinsky et al., 2013, Hongo & Yamano, 2013, Yara et al., 2011). In particular, rising sea surface temperatures have affected species distributions and diversity, with warm-adapted species expanding their ranges, while cold-adapted species are experiencing range contractions or shifts (Pinsky et al., 2013). For example, in Sekisei Lagoon, Okinawa, approximately 60-80% of corals became bleached during anomalously high temperatures in 2016, leading to a significant reduction in coral cover (Morita et al., 2020). Additionally, Beaugrand et al. (2023) reported a 24% decline in marine plankton abundance over the past 80 years, along with latitudinal and bathymetric shifts in their distribution. To better understand biological impacts of climate change, this study focused on nudibranchs.

Nudibranchs, belonging to the subclass Heterobranchia in the class Gastropoda of the phylum Mollusca, are a taxonomic group well adapted to a wide range of environments, from tropical to polar regions (Nakano, 2019; Gosliner et al., 2008). Approximately 6,000 species have been reported worldwide, with over 1,400 species documented in Japan alone (Kashio et al., 2021; Nakano, 2019). They are known for their diverse morphologies and ecological traits. By feeding on sponges, cnidarians, bryozoans, tunicates, and algae, nudibranchs help regulate populations of these organisms and contribute to maintenance of community structures (Gosliner et al., 2008). Recent studies have shown that nudibranchs respond sensitively to environmental changes, and they are increasingly considered bioindicators of biodiversity. For example, surveys along the eastern coast of Tottori Prefecture, Japan and surrounding marine areas have indicated that nudibranch species composition may serve as an indicator of benthic invertebrate communities influenced by the Tsushima Warm Current (Oota, 2021). In another case, nudibranchs have been recognized as effective biological indicators in environmental monitoring and assessment, owing to their sensitivity to changes in water quality, habitat conditions, and biodiversity (Chou et al., 2022; Atlas of Living Australia, 2021).

The nudibranch fauna is often documented through visual observations or photographs taken by diving instructors and recreational divers, but systematic studies of their species composition and community structure are limited, with most studies having been conducted only once (Andrimida, A. 2022; Kashio et al., 2021; Ota, 2021). The northwestern coast of Kyushu, Japan, faces diverse marine influences, including the East China Sea, the Genkai Sea, and the Tsushima Strait. Interactions among these elements foster rich biodiversity and complex ecosystems (Fujikura et al., 2010; Itsukushima & Kano, 2022). In this coastal region, two previous studies have investigated nudibranch community structures: one by Matsubayashi (1989) conducted during the 1960s–1980s, and another by Kawahara (unpublished) in 2003. In the present study, we focused on Tatsunokuchi and Akase in Nomozaki, located in northwestern Kyushu, Japan, near sites used in these earlier surveys. We conducted extensive visual surveys by SCUBA diving. By comparing species occurrence data obtained from the literature and our present study, we documented long-term changes in nudibranch species composition and biodiversity.

## 2. Materials and Methods

### 2.1. Historical faunal records

The survey conducted by Matsubayashi (1989) is estimated to have been carried out from the 1960s and 1980s, although the specific timing and frequency of sampling could not be documented. The study covered a wide geographic area, including offshore islands in Nagasaki Prefecture, Japan. To compare nudibranch fauna across decades, data from sites geographically close to the present study, (Mogi, Nomozaki, Kodatejin-iwa, Odatejin-iwa, Tanoko-jima, Takahama Ingeri-bana, and Meoto-iwa) were extracted (Fig. 1 and Supplementary Table S1). Nudibranchs were collected manually from various microhabitats, such as undersides and crevices of rocks, in tidal pools, overturned stones, larger tidal pools, and seaweed. Additional methods included sieving sand brought back from octopus pot fishing and collecting bycatch mollusks during shrimp trawl sorting operations. Species identification was based on morphological features, including radula structure, and followed a classification system for opisthobranchs. A full list of observed species is provided in Supplementary Table S1.

**Fig. 1.**
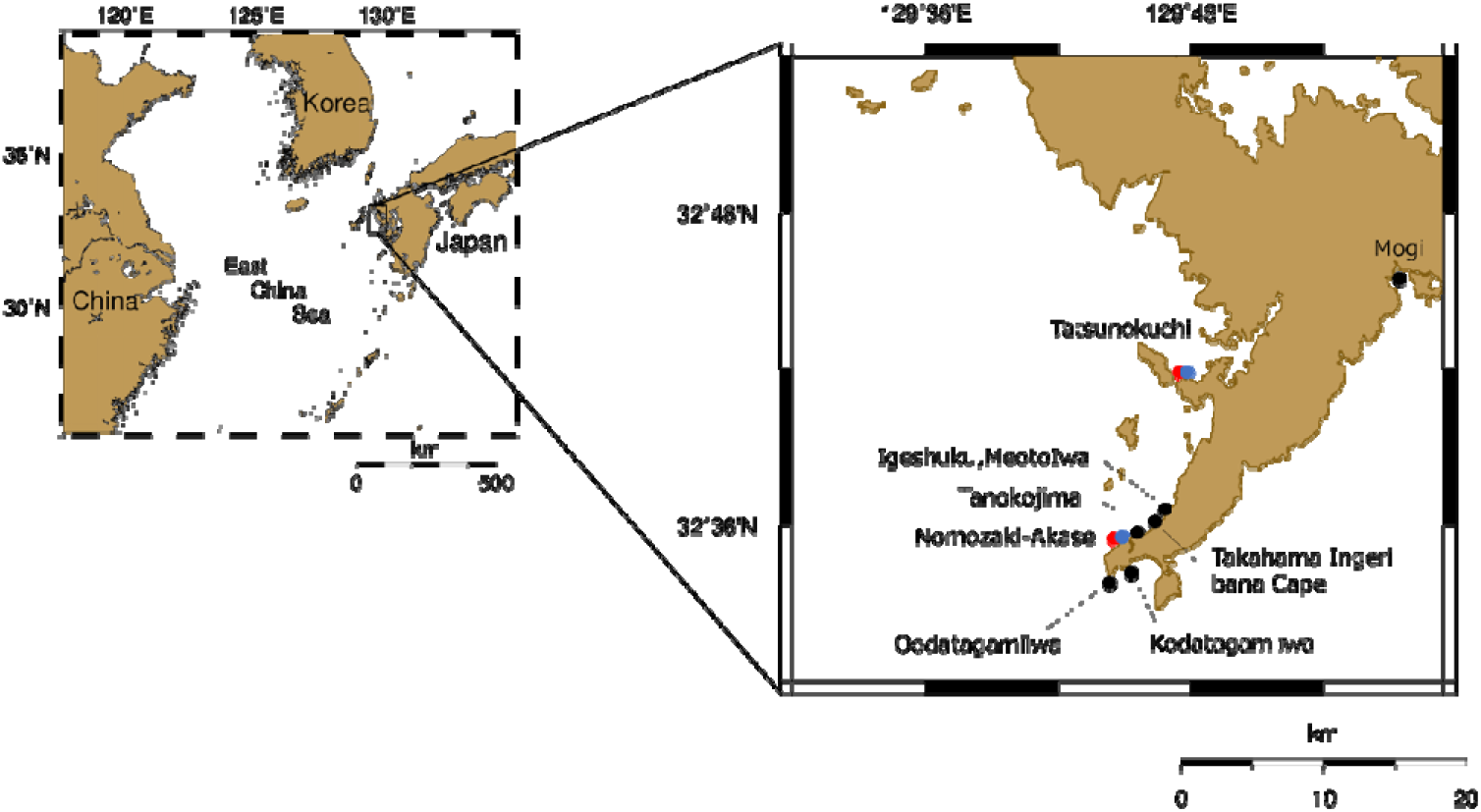
Survey sites of nudibranch assemblages. Black dots indicate sites surveyed by Matsubayashi (1989). Blue dots indicate sites surveyed by Kawahara (unpublished), and red dots indicate sites surveyed in the present study. Photographs of Kawahara (unpublished) and the present study sites (Tatsunokuchi and Nomozaki-Akase) are shown in Supplementary Fig. S1.

The survey conducted by Kawahara (unpublished) was compiled as a master’s thesis at the Faculty of Fisheries, Nagasaki University, but is not publicly available; thus, it is referred to here as unpublished. That survey was conducted from April 2001 to September 2003 using a combination of free diving, SCUBA diving, and intertidal collection. Although sampling was conducted at six sites in northwestern Kyushu, Japan, for temporal comparison in this study, we extracted data only from two of them (Tatsunokuchi and Nomozaki-Akase), due to the completeness of the dataset and their geographic correspondence with the present survey (Fig. 1 and Supplementary Table S1). Kawahara’s classification system followed Beesley, Ross, and Wells (1998), and species were identified based not only on external features, but also on internal anatomy such as the radula, digestive, and reproductive systems. A complete species list is provided in Supplementary Table S1.

### 2.2. SCUBA diving survey

To investigate the present composition of nudibranch assemblages, SCUBA diving surveys were conducted monthly from June 2023 to January 2024 at two coastal sites in northwestern Kyushu, Japan (Tatsunokuchi and Nomozaki Akase) (Supplementary Fig. S2). Surveys were carried out during daytime low tide at depths ranging from 3 to 15 meters. At each site, one or two dives lasting 40–80 min were performed. A free-search method was employed, targeting various microhabitats such as rocky reefs, boulders, and macroalgal beds. Two divers participated in each survey. One diver was responsible for photographing nudibranchs underwater using an OLYMPUS Tough TG-6 digital camera, while the other collected specimens or documented individuals based on photographs taken. Collected specimens were later examined under a stereomicroscope for detailed observation. Species identification was conducted based on morphological characteristics, referring to identification guides by Nakano (2019) and Gosliner et al. (2008). Individuals that could not be identified to species level were recorded as “cf. species name” or “sp.” In addition to biological data, environmental parameters such as water temperature, visibility, and substrate type were recorded immediately after each dive (Supplementary Table S2). Surveys were conducted for academic research purposes with official permission granted by the Nagasaki Prefectural Fisheries Division (Permit Nos. 5Gyoshin-Kyo-115 and -116). All dives were carried out under the supervision of a certified diving instructor for safety management.

### 2.3. Data analysis

Species diversity was compared using data collected by Kawahara (unpublished) from 2001 to 2003 and our survey data from 2023 to 2024. Data from Matsubayashi (1989) were excluded from diversity index calculations and comparative analyses due to the lack of individual counts. To evaluate species diversity, we used the number of observed species, the Shannon–Wiener diversity index (H), and Simpson’s diversity index (D).

The Shannon–Wiener index (H), which accounts for species evenness, was calculated as follows (Mittelbach & McGill, 2023):

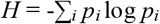

where *p*_*i*_ is the relative abundance of species *i*. Simpson’s diversity index (*D*), which emphasizes dominance, was calculated using the following formula (Mittelbach & McGill, 2023):

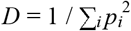

where *p*_*i*_ represents the relative abundance of species *i*.

To evaluate temporal changes in species composition, we calculated the relative abundance of each nudibranch species during each survey period (2001–2003 and 2023–2024), defined as the proportion of individuals of each species to the total number of individuals. Fisher’s exact tests were conducted to assess differences in species-specific proportions between the two periods. Resulting p-values were adjusted using the Benjamini–Hochberg procedure to control the false discovery rate (FDR). Species that showed statistically significant changes in composition (adjusted p < 0.05) were identified, resulting in identification of 14 species with significant shifts in relative abundance.

To compare temporal changes in the climatic preferences of nudibranch species, all recorded species were categorized into four climate affinity groups based on their primary distributional ranges: (1) tropical–subtropical, (2) temperate, (3) tropical–subtropical–temperate, and (4) subarctic–arctic. This classification scheme was adapted from the framework proposed by Grandéz et al. (2023) for nudibranchs, with modifications to better reflect distributional characteristics of species in Japanese coastal waters. In particular, species primarily distributed in northern Hokkaido and the Russian Far East were categorized as “subarctic–arctic.” The primary distribution of each species was determined based on the illustrated guide by Ono and Kato (2023). For species not listed in that guide, supplemental information was obtained from the user-submitted online database “World of Sea Slugs” (Nobuhiko, 2017). Species distributed mainly in the Indo-West Pacific, central Pacific, or tropical Pacific were classified as tropical–subtropical; those mainly found in the temperate zones of Japan, the Korean Peninsula, and Hong Kong were categorized as temperate. Species with wide distributions spanning multiple climatic zones, e.g., Indo-Pacific, Indo-West Pacific to temperate Japan, were assigned to the tropical–subtropical–temperate group. The same classification criteria were applied to species recorded in past surveys (Matsubayashi, 1989; Kawahara, unpublished). Species for which primary distribution ranges could not be verified due to lack of records in modern literature or databases were excluded from the climatic group analysis. A full list of species with their assigned climatic preferences is provided in Supplementary Tables S1 and S3.

To evaluate the similarity of species composition among the three survey periods (1960–1980, 2001–2003, and 2023–2024), we calculated the Jaccard Coefficient of Commonality (CC) (Kimoto and Takeda, 1989), defined as:

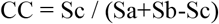

where Sa and Sb are the numbers of species recorded in each of the two periods, and Sc is the number of species common to both periods. A CC value of 0 indicates no species overlap between periods, whereas a value of 1 indicates complete overlap. In addition, we calculated the Sørensen–Dice Index (SDI) to further assess similarity in species composition:

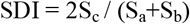

Like CC, SDI quantifies species overlap between two time periods, but places greater emphasis on shared species, typically yielding higher values than the Jaccard coefficient. Both indices were used to compare temporal shifts in nudibranch assemblages.

## 3. Results

In our SCUBA surveys conducted during 2023 and 2024, a total of 27 dives (equivalent to 25.45 hours) yielded 47 species and 304 individual nudibranchs (Tatsunokuchi: 30 species, 169 individuals; Nomozaki-Akase: 32 species, 135 individuals). In contrast, the 2001–2003 survey recorded 35 species (Tatsunokuchi: 25 species, 313 individuals; Nomozaki-Akase: 21 species, 189 individuals) (Fig. 2a). The Shannon–Wiener diversity index (*H*) was 2.21 in 2001–2003 and 3.10 in 2023–2024, representing a 40.3% increase in recent years (Fig. 2b). Similarly, the Simpson diversity index (*D*) increased from 0.76 to 0.93, corresponding to a 22.4% rise (Fig. 2c). By either measure, species diversity has increased significantly over the past two decades.

**Fig. 2.**
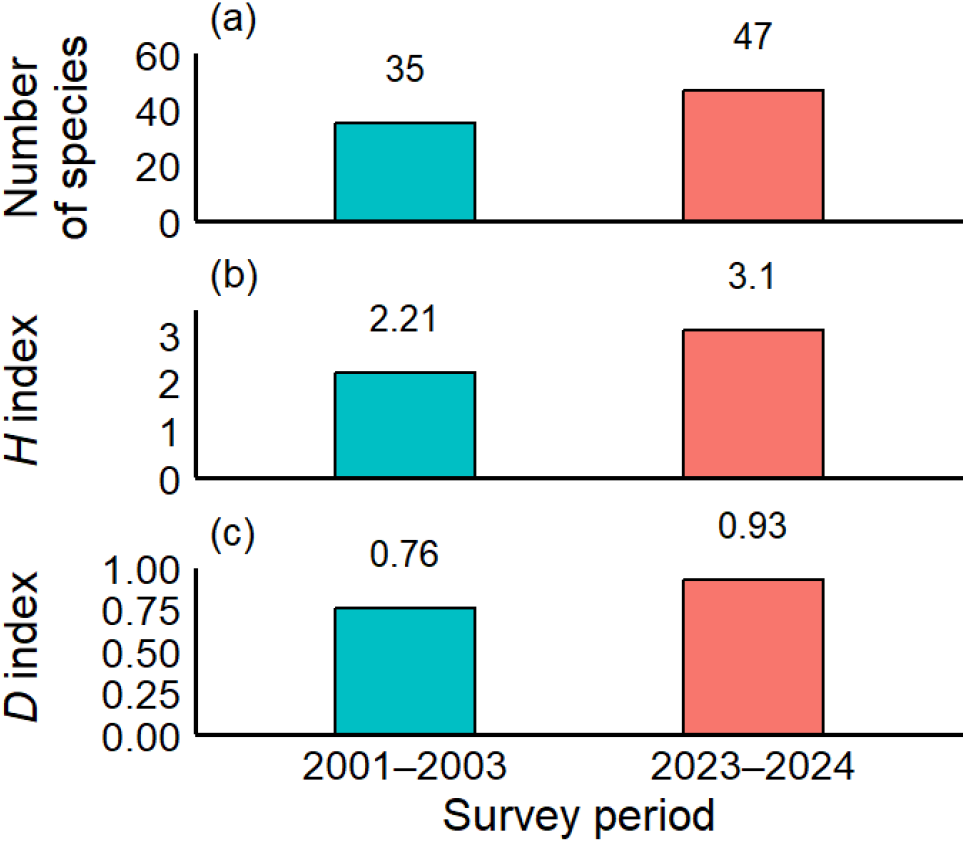
Comparison of nudibranch assemblages in the past (2001–2003) and the present (2023–2024). (a) Numbers of species, (b) Shannon-Wiener diversity index (*H*), and (c) Simpson diversity index (*D*). Both surveys were conducted at Tatsunokuchi and Nomozaki-Akase on the west coast of Kyushu, Japan.

Community structure, based on relative abundances of individual species, showed a clear shift between 2001–2003 and 2023–2024 (Fig. 3). In the earlier period (2001–2003), a few dominant species, such as *Aplysia kurodai* and *Aplysia oculifera*, accounted for a large proportion of the community, with *Aplysia kurodai* alone comprising approximately 45% of all individuals. In contrast, in the recent period (2023–2024), the relative abundance of these dominant species declined markedly. Instead, species such as *Tritoniopsis elegans, Doriprismatica atromarginata, Goniobranchus orientalis, Goniobranchus sinensis, and Goniobranchus tinctorius* increased in proportion, indicating a shift toward a more diverse community. These compositional shifts were further supported by statistical analyses, which identified 15 species exhibiting significant changes in relative abundance between the 2001–2003 and 2023–2024 periods (Fig. 4).

**Fig. 3.**
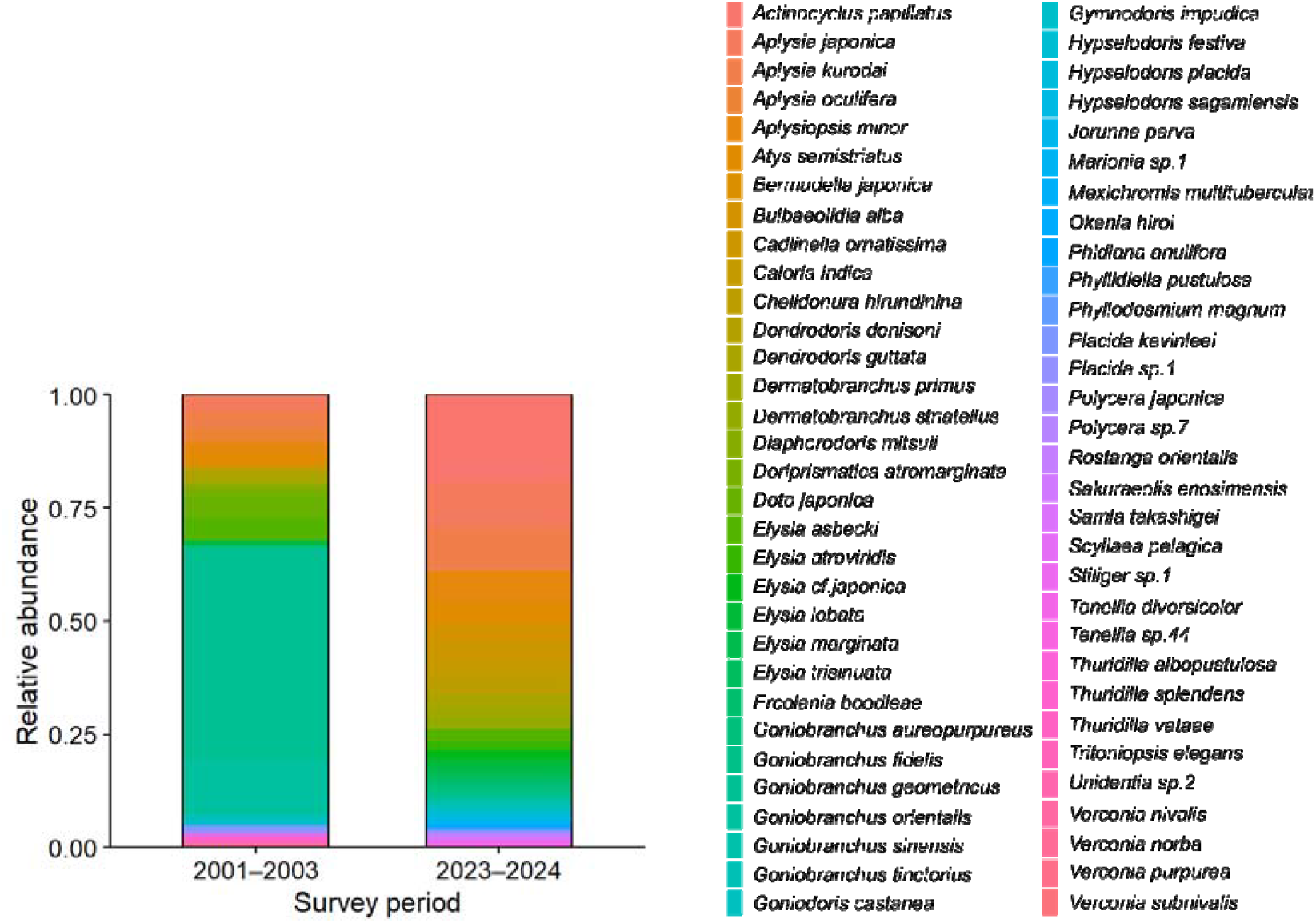
Comparative heatmap of the relative abundance of nudibranch assemblages in the past (2001–2003) and present (2023–2024). Relative abundance was calculated as the proportion of individuals of each species to the total number of individuals observed during each period. Both surveys were conducted at Tatsunokuchi and Nomozaki-Akase on the west coast of Kyushu, Japan.

**Fig. 4.**
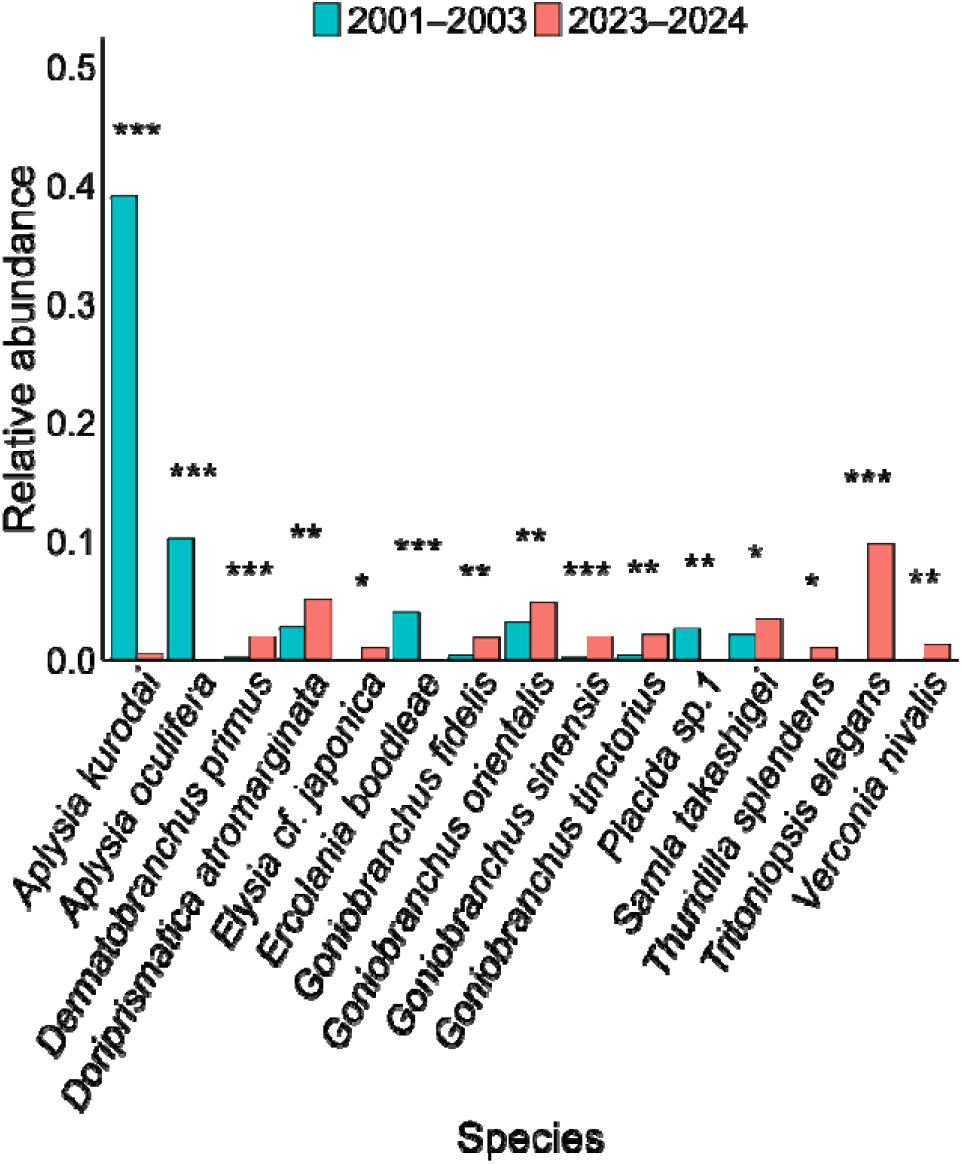
Comparison of relative abundances of 15 nudibranch species that showed statistically significant differences between the past (2001–2003) and present (2023–2024). Asterisks indicate significance levels: \**p* < 0.05, \*\**p* < 0.01, \*\*\**p* < 0.001. Both surveys were conducted at Tatsunokuchi and Nomozaki-Akase on the northwestern coast of Kyushu, Japan.

Comparison of species composition by climatic preferences revealed that among the 47 species observed in the present study (2023–2024), 55.3% (26 species) were tropical–subtropical, 19.2% (9 species) were tropical–subtropical–temperate, and 25.5% (12 species) were temperate. In contrast, among the 79 species recorded between 1960 and 1980 (Matsubayashi, 1989), 43.0% (34 species) were tropical–subtropical, 22.8% (18 species) were tropical–subtropical–temperate, 31.6% (25 species) were temperate, and 2.5% (2 species) were subarctic–arctic. Similarly, among the 35 species recorded approximately 20 years ago (Kawahara, unpublished), 42.9% (15 species) were tropical–subtropical, 25.7% (9 species) were tropical–subtropical–temperate, 28.6% (10 species) were temperate, and 2.9% (1 species) were subarctic–arctic. These results indicate increasing proportions of tropical–subtropical species and decreasing proportions of temperate species (Fig. 5). The number of opisthobranch species observed in each time period was 47 species in 2023–2024, 35 species in 2001–2003, and 79 species in 1960–1980. Among the three periods, the lowest similarity in species composition was found between 2023–2024 and 1973–1989 (Jaccard Coefficient [CC]: 0.105; Sørensen–Dice Index [SDI]: 0.190) (Table 1). In contrast, the similarity between 2001–2003 and 1973–1989 was relatively higher (CC: 0.281; SDI: 0.439) (Table 1).

**Table 1.** Numbers of shared species, Jaccard Coefficient (CC), and Sørensen–Dice Index (SDI) for opisthobranch species observed in different survey periods along the northwestern coast of Kyushu, Japan.

| Comparison Period | Number of Shared Species | Jaccard Coefficient | Sorensen-Dice Index |
| --- | --- | --- | --- |
| 1960-1980×2001-2003 | 18 | 0.188 | 0.316 |
| 2001-2003×2023-2024 | 18 | 0.281 | 0.439 |
| 1960-1980×2023-2024 | 12 | 0.105 | 0.191 |

**Fig. 5.**
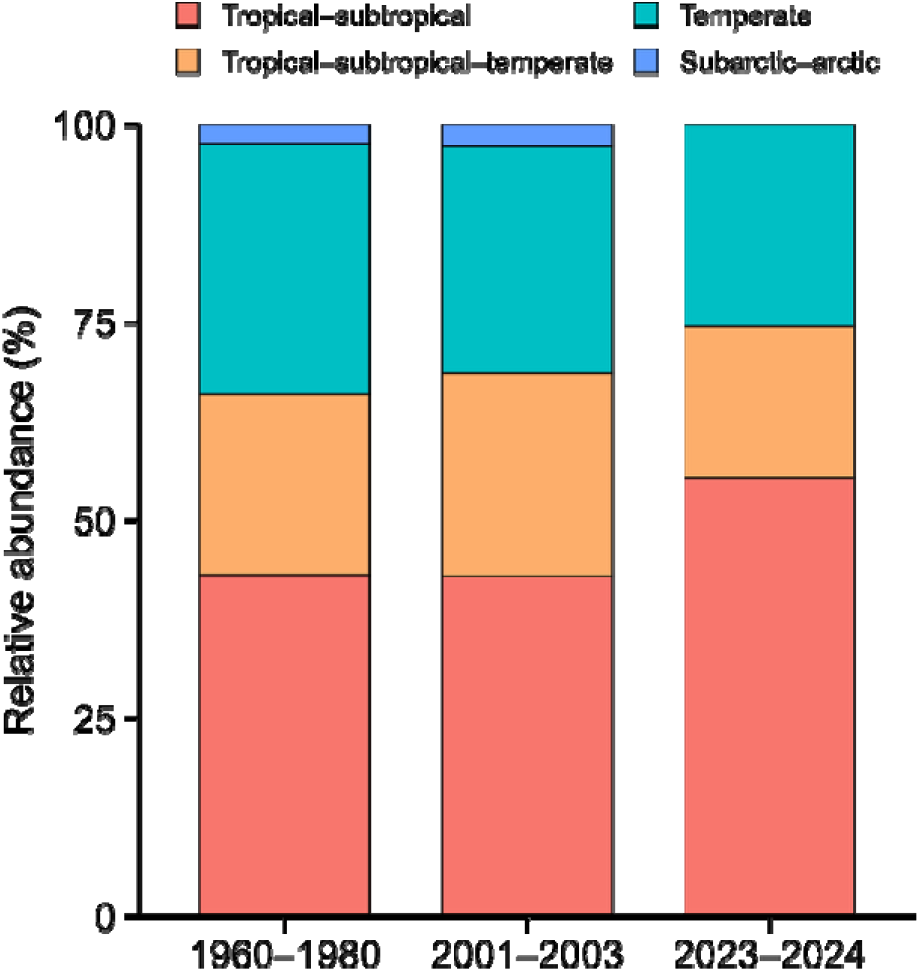
Comparison of climatic preferences of observed nudibranch species during different survey periods. In the most recent period, species belonging to the subarctic–arctic group were no longer detected, while the proportion of tropical–subtropical species increased. All surveys were conducted along the northwestern coast of Kyushu, Japan.

## 4. Discussion

The sea and its nudibranch fauna have undergone dramatic changes over the past 50 years, almost appearing as entirely different worlds. In this study, we quantitatively and qualitatively compared nudibranch communities across three time periods (1960–1980, 2001–2003, and 2023–2024) to reveal long-term shifts in community structure. Compared to 20 years ago, diversity indices (Shannon–Wiener diversity index (*H*) and Simpson diversity index (*D*)) have increased, and statistically significant changes in species composition ratios were observed in 15 species, indicating dynamic alterations at the species level (Figs. 2 and 4). Heatmap analyses revealed clear shifts in community structure, with distinct differences in species composition between time periods (Fig. 3). Similarity analyses using Jaccard Coefficient of Commonality (CC) and Sørensen–Dice Index (SDI) showed the greatest community dissimilarity between the 1960–1980 and 2023–2024 periods, suggesting a major community turnover. Notably, approximately 70% of species newly observed in the 2023–2024 survey were of tropical–subtropical origin, suggesting an ongoing shift in species distributions likely driven by climate change. Despite limitations in historical survey methods and records, this study represents the first comprehensive attempt to assess long-term changes in marine communities using nudibranchs as bioindicators.

Why has the composition of nudibranch assemblages changed so dramatically? The most plausible explanation lies in recent climate change, particularly the rise in seawater temperature. In fact, long-term sea surface temperature (SST) anomalies in the study area have shown a marked and persistent warming trend, especially from the late 2010s into the 2020s (Fig. 6). Since the 1980s, SSTs have steadily increased, with strong positive anomalies evident in recent years. Such thermal changes drive poleward range expansions of tropical and subtropical species (Molinos et al., 2022; Poloczanska et al., 2013). This pattern is clearly reflected in our data. Of the 47 nudibranch species recorded in 2023–2024, 55.3% were classified as tropical–subtropical species, substantially higher than proportions observed in 1960-1980 (43.0%) and 2001–2003 (40.0%). Shallow-water nudibranchs are particularly sensitive to water temperature and exhibit rapid shifts in occurrence patterns in response to thermal variability (Sanford et al., 2016). In addition to temperature, other environmental factors likely contribute to community restructuring, including fluctuations in food availability, e.g., algae and encrusting organisms, changes in benthic substrate composition, and anthropogenic impacts such as coastal development (Enright et al., 2021; Herbert-Read et al., 2022). These factors act synergistically, suggesting that the recent reorganization of nudibranch assemblages is the result of a complex interplay between abiotic environmental changes and biotic interactions.

**Fig. 6.**
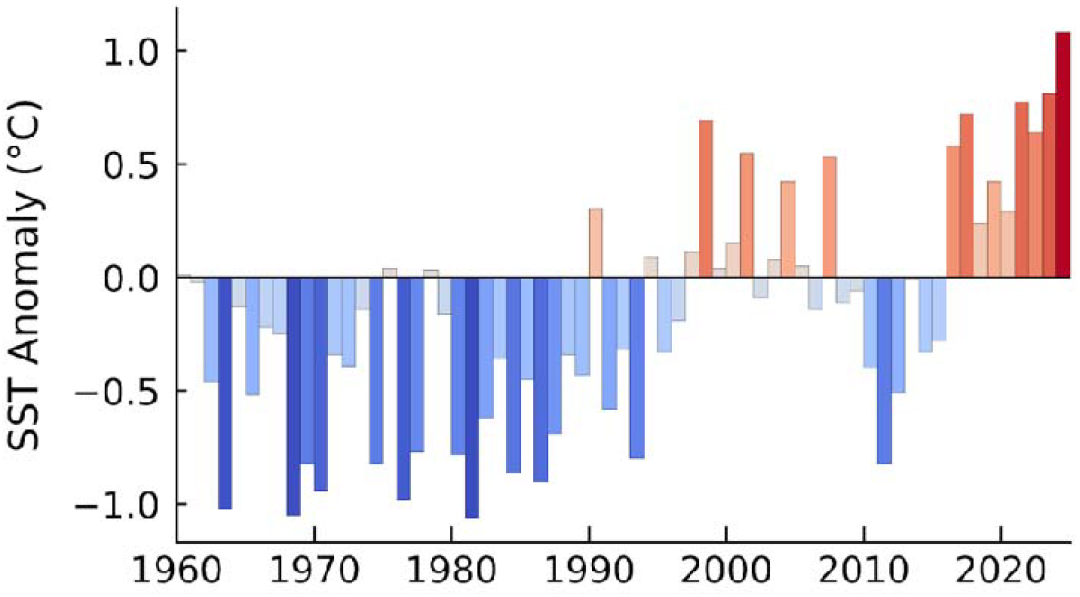
Annual sea surface temperature (SST) anomalies in the study region from 1960 to 2024. Each bar represents the deviation (in °C) from the 1991–2020 climatological average. Red bars indicate warmer-than-average years, while blue bars indicate cooler-than-average years. The height of each bar corresponds to the magnitude of the anomaly.

An important observation in this study is the emergence of multiple nudibranch species never recorded in this region during any previous surveys. Compared with earlier reports (Matsubayashi, 1989; Kawahara, unpublished), 24 species were newly documented along the northwest coast of Kyushu, Japan during the 2023–2024 surveys (Table 2). Notably, 17 of these new records (approximately 71%) were tropical to subtropical species, suggesting that populations originating from offshore waters during warm seasons may be establishing themselves in the coastal zone (Sanford et al., 2019; Sorte et al., 2010). Such poleward range expansions and climate-driven shifts in biodiversity have been reported globally (Poloczanska et al., 2013; Burrows et al., 2014), and the present findings suggest that similar community-level changes are already underway in this region as part of broader warming-driven biogeographic rearrangements (Dornelas et al., 2023; Pinsky et al., 2020). The arrival of previously unrecorded species may lead to novel competitive and trophic interactions, potentially disrupting existing ecological balances (Sasaki et al., 2024; Zarco□Perello & Wernberg, 2020). Newly observed species likely include undescribed or rare taxa, indicating that ecological changes may not be fully captured by conventional taxonomic or ecosystem assessment frameworks (Rogers et al., 2023). This lag in recognition poses challenges for biodiversity monitoring and conservation, emphasizing the urgent need for accurate species identification and systematic data accumulation (Ma et al., 2024, Rout et al., 2022). While the appearance of new species may superficially suggest increased biodiversity, it often accompanies a reorganization of local communities, including shifts in dominant species or disappearance of previously common taxa, which may trigger ecosystem instability and negative feedback loops (Pecl et al., 2017). These findings underscore the necessity of understanding climate-driven responses not only in terms of tropical species expansion, but also in the context of local extinctions of cold-affinity species, calling for a more holistic view of climate-induced ecological transformation.

**Table 2.**
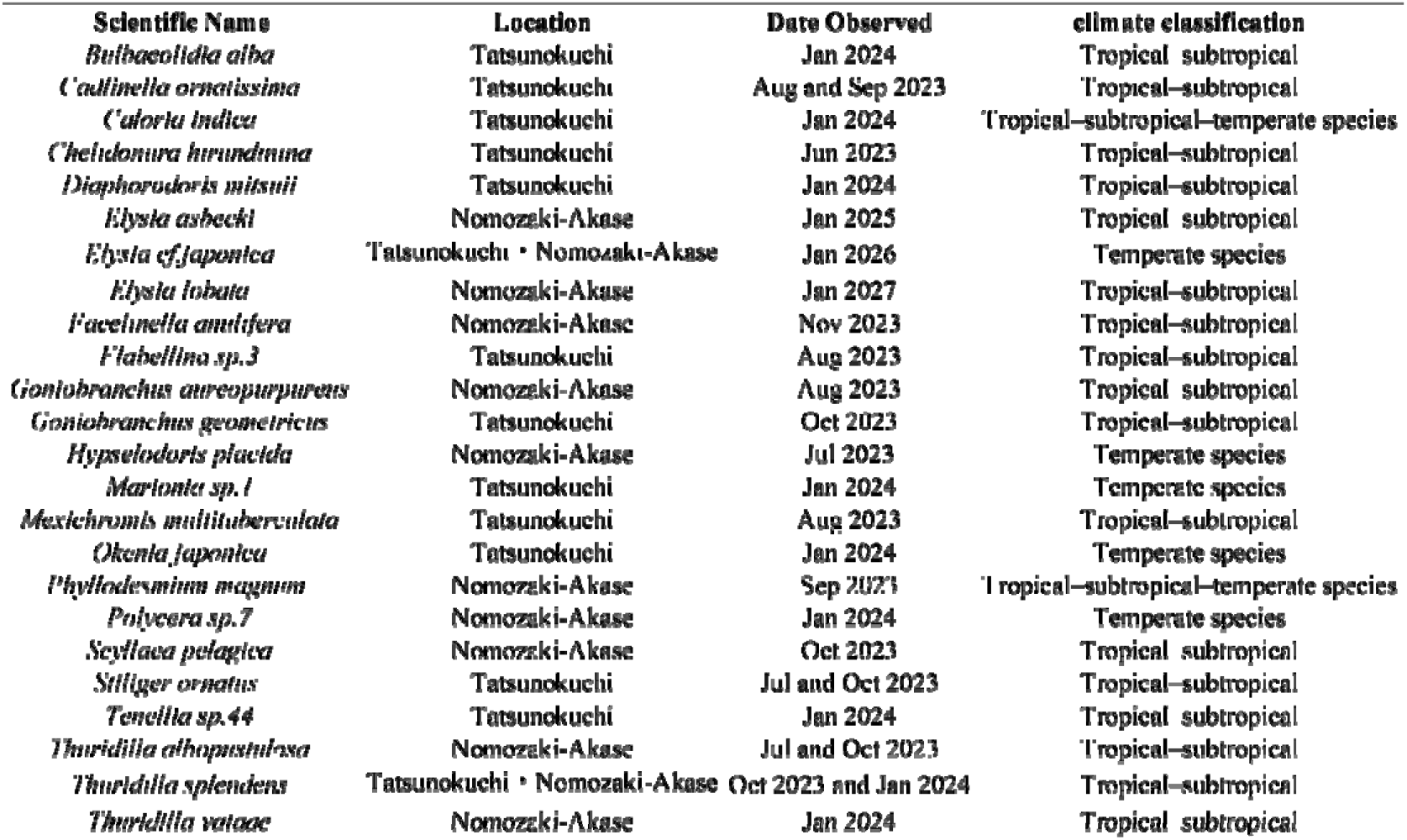
List of 24 opisthobranch species newly recorded in coastal waters of northwestern Kyushu during the 2023–2024 survey. Species not documented in previous literature (Matsubayashi, 1989; Kawahara, unpublished) are considered newly recorded for the region. Names such as sp.1 are provisional identifiers used to distinguish undescribed or unidentified species in the same genus.

This study has certain limitations, particularly in quantitative comparisons with historical data from 50 years ago due to missing individual counts and unspecified survey periods in earlier datasets. Moreover, differences in survey methods and the limited number of sampling sites must also be acknowledged as constraints. The nature of scuba-based surveys introduces additional variability due to diver experience and sea conditions. Indeed, previous studies have reported that observer expertise and experience can influence species detection rates and abundance estimates in underwater visual censuses (Coleman et al., 2013). Sea surface temperature rise due to global warming is accelerating, and its consequences remain largely unknown. How will the ocean change in future decades? To better answer this question, more quantitative and standardized monitoring approaches are essential. Incorporating methods such as line transect surveys can help reduce observer bias and improve data consistency. Additionally, the use of environmental DNA (eDNA) and image analysis technologies can enable more extensive and objective monitoring of biodiversity. Furthermore, collecting information on the age structure and reproductive status of observed individuals will be valuable in understanding processes of range expansion and local establishment. Continued quantitative, long-term monitoring of coastal ecosystems is vital for developing effective conservation strategies and improving predictions of future ecological shifts.

## Acknowledgements

We express our sincere gratitude to Mr. Koji Sugisaki of the diving shop *Smilers* (https://smilers.jp/) for ensuring the safety of our diving surveys. We also extend our deep appreciation to members of the Nomozaki Sanwa Fisheries Cooperative and the Seihi Southern Fisheries Cooperative for granting us permission to conduct the underwater surveys. Finally, we thank Steven D. Aird, Technical Editor (https://www.sda-technical-editor.org/), and journal editors and anonymous reviewers for their valuable comments and suggestions that greatly improved the quality of the manuscript. This work was supported by the Sasakawa Scientific Research Grant from the Japan Science Society (Grant Number: 2024-4058).

## Competing Interests

The authors have no competing interests to declare.

## Ethics Approval

The research required no permit approvals.

## Author Contributions

RK Investigation, Writing – original draft, Visualization. ST Investigation, Writing – review & editing. MY Conceptualization, Methodology, Investigation, Writing – original draft, Supervision, Funding acquisition, Visualization.

## Supporting Information

Additional supporting information can be found online in the Supporting Information.

## Data Availability

Data has been provided as Supplementary Material.

## Notes

### Competing Interest Statement

The authors have declared no competing interest.

