## Supplementary Information for "A different world: temporal changes in nudibranch community structure over a half-century"

**Table S1** Summary of nudibranch data extracted from the literature. (a) Matsubayashi (1989); (b) Kawahara (unpublished). Climatic distribution categories, e.g., tropical-subtropical, warm-temperate, were assigned to each species in this study.

(a) From Matsubayashi (1989).

| **Site** | **Order** | **Suborder** | **Superfamily** | **Family** | **Subfamily** | **Genus** | **Species** | **Climate classification** |
| --- | --- | --- | --- | --- | --- | --- | --- | --- |
| Mogi | Pleurobranchida |  |  | Pleurobranchoidae |  | Pleurobranchaea | *Pleurobranchaea maculata*  (Quoy&Gaimard,1832) | Tropical–subtropical species |
|  | Cladobranchia |  |  | Arminidae |  | Armina | *Armina* sp.2 | Temperate species |
|  | Nudibranchia | Doridacea(Phanerobranchia) |  | Plolyceridae | Kalininae | Kalininae | *Kalinga ornata*  (Alder&Hancock,1864) | Tropical–subtropical–temperate species |
| Nomozaki | Acteonimorpha |  |  | Acteonidae |  | Punctacteon | *Punctacteon fabreanus*  (Crosse,1874) | Tropical–subtropical species |
|  | Cephalaspidea |  |  | Tornatinidae |  | Genus Acteocina | *Acteocina exilis* (Dunker, 1860) | Temperate species |
|  | Aplysiida |  |  | Aplysiidae |  | Aplysia | *Aplysia japonica*  (G.B.Sowerby Ⅱ,1869) | Temperate species |
|  | Umbraculoidea |  |  | Umbraculidae |  | Umbraculum | *Umbraculum umbraculum*  (Lightfoot,1786) | Tropical–subtropical–temperate species |
|  | Nudibranchia | Doridacea(Phanerobranchia) |  | Polyceridae | Triophinae | Kaloplocamus | *Kaloplocamus acutus* Baba,1955 | Tropical–subtropical species |
|  | Nudibranchia | Doridacea(Phanerobranchia) |  | Polyceridae | Triophinae | Crimora | *Crimora lutea* Baba,1949 | Tropical–subtropical–temperate species |
|  | Nudibranchia | Doridacea(Cryptobranchia) |  | Chromodorididae |  | Goniobranchus | *Goniobranchus tumuliferus*  (Collingwood,1881) | Tropical–subtropical species |
|  | Cladobranchia |  |  | Arminidae |  | Dermatobranchus | *Dermatobranchus ornatus*  (Bergh,1874) | Tropical–subtropical species |
| Kodatagamiiwa | Aplysiida |  |  | Aplysiidae |  | Aplysia | *Aplysia japonica*  (G.B.Sowerby Ⅱ,1869) | Temperate species |
|  | Aplysiida |  |  | Aplysiidae |  | Aplysia | *Aplysia kurodai*(Baba,1937) | Tropical–subtropical species |
|  | Aplysiida |  |  | Aplysiidae |  | Dolabella | *Dolabella auricularia*  (Lightfoott,1786) | Tropical–subtropical species |
|  | Sacoglossa |  |  | Limapontiidae |  | Placida | *Placida* sp.1 | Temperate species |
|  | Sacoglossa |  |  | Hermaeidae |  | Aplysiopsis | *Aplysiopsis nigra*(Baba,1949) | Temperate species |
|  | Sacoglossa |  |  | Plakobranchidae |  | Elysia | *Elysia atroviridis* Baba,1955 | Temperate species |
|  | Nudibranchia | Doridacea(Phanerobranchia) |  | Gymnodoridae |  | Gymnodoris | *Gymnodoris citrina*  (Bergh,1877) | Tropical–subtropical–temperate species |
|  | Nudibranchia | Doridacea(Phanerobranchia) |  | Okadaiidae |  | Vayssierea | *Vayssierea felis*  (Collingwood,1881) | Tropical–subtropical species |
|  | Nudibranchia | Doridacea(Cryptobranchia) |  | Chromodorididae |  | Goniobranchus | *Goniobranchus orientails*  (Rudman,1983) | Tropical–subtropical species |
|  | Nudibranchia | Doridacea(Cryptobranchia) |  | Chromodorididae |  | Hypselodoris | *Hypselodoris festiva*(A.Adams,1861) | Temperate species |
|  | Nudibranchia | Doridacea(Cryptobranchia) |  | Chromodorididae |  | Hypselodoris | *Hypselodoris maritima*  (Baba,1949) | Tropical–subtropical species |
|  | Nudibranchia | Doridacea(Cryptobranchia) |  | Actinocyclidae |  | Actinocyclus | *Actinocyclus papillatus*  (Bergh,1878) | Tropical–subtropical species |
|  | Nudibranchia | Doridacea(Cryptobranchia) |  | Discodorididae |  | Rostanga | *Rostanga orientalis* Rudman&Avern,1989 | Temperate species |
|  | Nudibranchia | Doridacea(Cryptobranchia) |  | Dorididae |  | Doris | *Doris pecten* Collingwood,1881 | Tropical–subtropical species |
|  | Nudibranchia | Doridacea(Cryptobranchia) |  | Discodorididae |  | Discodoris | *Discodoris lilacina*(Gould,1852) | Tropical–subtropical species |
|  | Nudibranchia | Doridacea(Porostomata) |  | Dendorodorididae |  | Dendrodoris | *Dendrodoris arborescens*  (Collingwood,1881) | Tropical–subtropical–temperate species |
|  | Cladobranchia |  |  | Arminidae |  | Dermatobranchus | *Dermatobranchus otome* Baba,1992 | Temperate species |
|  | Cladobranchia |  | Fionidea | Fionidae |  | Tenellia | *Tenellia ornata*(Baba,1937) | Tropical–subtropical species |
| Oodatagamiiwa | Aplysiida |  |  | Aplysiidae |  | Aplysia | *Aplysia japonica*  (G.B.Sowerby Ⅱ,1869) | Temperate species |
|  | Aplysiida |  |  | Aplysiidae |  | Aplysia | *Aplysia kurodai*(Baba,1937) | Tropical–subtropical species |
|  | Aplysiida |  |  | Aplysiidae |  | Dolabella | *Dolabella auricularia*  (Lightfoott,1786) | Tropical–subtropical species |
|  | Sacoglossa |  |  | Limapontiidae |  | Placida | *Placida* sp.1 | Temperate species |
|  | Sacoglossa |  |  | Limapontiidae |  | Ercolania | *Ercolania boodleae*(Baba,1938) | Subarctic–arctic species |
|  | Sacoglossa |  |  | Plakobranchidae |  | Elysia | *Elysia atroviridis* Baba,1955 | Temperate species |
|  | Sacoglossa |  |  | Elysiidae |  | Elysia | *Elysia sugashimae* Baba, 1955 | Temperate species |
|  | Nudibranchia | Doridacea(Phanerobranchia) |  | Goniodorididae |  | Goniodoris | *Goniodoris joubini* Risbec,1928 | Tropical–subtropical species |
|  | Nudibranchia | Doridacea(Phanerobranchia) |  | Okadaiidae |  | Vayssierea | *Vayssierea felis*  (Collingwood,1881) | Tropical–subtropical species |
|  | Nudibranchia | Doridacea(Cryptobranchia) |  | Chromodorididae |  | Goniobranchus | *Goniobranchus orientails*  (Rudman,1983) | Tropical–subtropical species |
|  | Nudibranchia | Doridacea(Cryptobranchia) |  | Chromodorididae |  | Hypselodoris | *Hypselodoris festiva*  (A.Adams,1861) | Temperate species |
|  | Nudibranchia | Doridacea(Porostomata) |  | Dendorodorididae |  | Doriopsilla | *Doriopsilla miniata*  (Alder&Hancock,1864) | Tropical–subtropical species |
|  | Cladobranchia |  |  | Arminidae |  | Dermatobranchus | *Dermatobranchus otome* Baba,1992 | Temperate species |
|  | Cladobranchia |  | Fionidea | Fionidae |  | Tenellia | *Tenellia anulata*(Baba,1949) | Tropical–subtropical species |
| Tanokojima | Cephalaspidea |  |  | Haminoeidae |  | Atys | *Atys semistriatus* Pease,1860 | Tropical–subtropical–temperate species |
|  | Cephalaspidea |  |  | Gastropteridade |  | Siphopteron | *Siphopteron brunneomarginatum*  (Carlson&Hoff,1974) | Tropical–subtropical–temperate species |
|  | Aplysiida |  |  | Aplysiidae |  | Dolabella | *Dolabella auricularia*  (Lightfoott,1786) | Tropical–subtropical species |
|  | Aplysiida |  |  | Aplysiidae |  | Aplysia | *Aplysia japonica*  (G.B.Sowerby Ⅱ,1869) | Temperate species |
|  | Aplysiida |  |  | Aplysiidae |  | Aplysia | *Aplysia kurodai*(Baba,1937) | Tropical–subtropical species |
|  | Aplysiida |  |  | Aplysiidae |  | Petalifera | *Petalifera punctulata*  (Tapparone-Canefri,1874) | Tropical–subtropical–temperate species |
|  | Aplysiida |  |  | Aplysiidae |  | Dolabrifera | *Dolabrifera dolabrifera*  (Cuvier,1817) | Tropical–subtropical species |
|  | Sacoglossa |  |  | Plakobranchidae |  | Elysia | *Elysia amakusana* Baba,1955 | Tropical–subtropical species |
|  | Sacoglossa |  |  | Plakobranchidae |  | Elysia | *Elysia obtusa* Baba,1938 | Tropical–subtropical–temperate species |
|  | Sacoglossa |  |  | Plakobranchidae |  | Elysia | *Elysia atroviridis* Baba,1955 | Temperate species |
|  | Sacoglossa |  |  | Plakobranchidae |  | Elysia | *Elysia nigrocapitata* Baba, 1957 | Temperate species |
|  | Sacoglossa |  |  | Plakobranchidae |  | Elysia | *Elysia trisinuata* Baba,1949 | Tropical–subtropical species |
|  | Sacoglossa |  |  | Elysiidae |  | Elysia | *Elysia sugashimae* Baba, 1955 | Temperate species |
|  | Sacoglossa |  |  | Elysiidae |  | Elysia | *Elysia marginata*(Pease,1871) | Tropical–subtropical–temperate species |
|  | Sacoglossa |  |  | Limapontiidae |  | Placida | *Placida kevinleei* McCarthy,Krug&Valdes,2017 | Tropical–subtropical–temperate species |
|  | Sacoglossa |  |  | Limapontiidae |  | Placida | *Placida* sp.1 | Temperate species |
|  | Sacoglossa |  |  | Limapontiidae |  | Ercolania | *Ercolania boodleae*  (Baba,1938) | Subarctic–arctic species |
|  | Sacoglossa |  |  | Hermaeidae |  | Aplysiopsis | *Aplysiopsis nigra*(Baba,1949) | Temperate species |
|  | Pleurobranchida |  |  | Pleurobranchoidae |  | Berthella | *Berthella stellata*(Risso,1826) | Tropical–subtropical–temperate species |
|  | Pleurobranchida |  |  | Pleurobranchoidae |  | Pleurobranchaea | *Pleurobranchaea maculata*  (Quoy&Gaimard,1832) | Tropical–subtropical species |
|  | Nudibranchia | Doridacea(Phanerobranchia) |  | Polyceridae | Polycerinae | Polycera | *Polycera fujitai* Baba,1937 | Tropical–subtropical species |
|  | Nudibranchia | Doridacea(Phanerobranchia) |  | Polyceridae | Polycerinae | Palio | *Palio amakusana* Baba,1960 | Temperate species |
|  | Nudibranchia | Doridacea(Phanerobranchia) |  | Gymnodoridae |  | Gymnodoris | *Gymnodoris impudica*  (Ruppell&Leuckart,1828) | Tropical–subtropical species |
|  | Nudibranchia | Doridacea(Phanerobranchia) |  | Gymnodoridae |  | Gymnodoris | *Gymnodoris subornata* Baba,1960 | Temperate species |
|  | Nudibranchia | Doridacea(Phanerobranchia) |  | Gymnodoridae |  | Gymnodoris | *Gymnodoris citrina*  (Bergh,1877) | Temperate species |
|  | Nudibranchia | Doridacea(Phanerobranchia) |  | Polyceridae | Triophinae | Kaloplocamus | *Kaloplocamus ramosus*  (Cantraine,1835) | Tropical–subtropical–temperate species |
|  | Nudibranchia | Doridacea(Phanerobranchia) |  | Polyceridae | Triophinae | Kaloplocamus | *Kaloplocamus acutus* Baba,1955 | Tropical–subtropical species |
|  | Nudibranchia | Doridacea(Phanerobranchia) |  | Goniodorididae |  | Okenia | *Okenia distinca* Baba,1940 | Temperate species |
|  | Nudibranchia | Doridacea(Phanerobranchia) |  | Goniodorididae |  | Goniodoris | *Goniodoris joubini* Risbec,1928 | Tropical–subtropical species |
|  | Nudibranchia | Doridacea(Phanerobranchia) |  | Goniodorididae |  | Goniodoris | *Goniodoris castanea* Alder&Hancock,1845 | Tropical–subtropical–temperate species |
|  | Nudibranchia | Doridacea(Phanerobranchia) |  | Goniodorididae |  | Okenia | *Okenia hiroi*(Baba,1938) | Tropical–subtropical species |
|  | Nudibranchia | Doridacea(Phanerobranchia) |  | Okadaiidae |  | Vayssierea | *Vayssierea felis*  (Collingwood,1881) | Tropical–subtropical species |
|  | Nudibranchia | Doridacea(Cryptobranchia) |  | Chromodorididae |  | Goniobranchus | *Goniobranchus orientails*  (Rudman,1983) | Tropical–subtropical species |
|  | Nudibranchia | Doridacea(Cryptobranchia) |  | Chromodorididae |  | Hypselodoris | *Hypselodoris festiva*  (A.Adams,1861) | Temperate species |
|  | Nudibranchia | Doridacea(Cryptobranchia) |  | Chromodorididae |  | Goniobranchus | *Goniobranchus tinctorius*  (Ruppell&Leuckart,1830) | Tropical–subtropical species |
|  | Nudibranchia | Doridacea(Cryptobranchia) |  | Chromodorididae |  | Verconia | *Verconia nivalis*(Baba,1937) | Temperate species |
|  | Nudibranchia | Doridacea(Cryptobranchia) |  | Chromodorididae |  | Goniobranchus | *Goniobranchus tumuliferus*  (Collingwood,1881) | Tropical–subtropical species |
|  | Nudibranchia | Doridacea(Cryptobranchia) |  | Discodorididae |  | Rostanga | *Rostanga orientalis* Rudman&Avern,1989 | Temperate species |
|  | Nudibranchia | Doridacea(Cryptobranchia) |  | Dorididae |  | Doris | *Doris pecten* Collingwood,1881 | Tropical–subtropical species |
|  | Nudibranchia | Doridacea(Cryptobranchia) |  | Discodorididae |  | Discodoris | *Discodoris lilacina*(Gould,1852) | Tropical–subtropical species |
|  | Nudibranchia | Doridacea(Cryptobranchia) |  | Discodorididae |  | Jorunna | *Jorunna parva*(Baba,1938) | Tropical–subtropical species |
|  | Nudibranchia | Doridacea(Cryptobranchia) |  | Discodorididae |  | Homoiodoris | *Homoiodoris japonica* Bergh,1881 | Temperate species |
|  | Nudibranchia | Doridacea(Cryptobranchia) |  | Discodorididae |  | Platydoris | *Platydoris ellioti*  (Alder&Hancock,1864) | Tropical–subtropical species |
|  | Nudibranchia | Doridacea(Porostomata) |  | Dendorodorididae |  | Dendrodoris | *Dendrodoris arborescens*  (Collingwood,1881) | Tropical–subtropical–temperate species |
|  | Nudibranchia | Doridacea(Porostomata) |  | Dendorodorididae |  | Dendrodoris | *Dendrodoris fumata*  (Ruppell&Leuckart,1830) | Tropical–subtropical–temperate species |
|  | Nudibranchia | Doridacea(Porostomata) |  | Dendorodorididae |  | Dendrodoris | *Dendrodoris denisoni*  (Angas,1864) | Tropical–subtropical–temperate species |
|  | Nudibranchia | Doridacea(Porostomata) |  | Dendorodorididae |  | Dendrodoris | *Dendrodoris guttata*  (Odhner,1917) | Tropical–subtropical species |
|  | Nudibranchia | Cladobranchia |  | Arminidae |  | Dermatobranchus | *Dermatobranchus otome* Baba,1992 | Temperate species |
|  | Nudibranchia | Cladobranchia |  | Arminidae |  | Dermatobranchus | *Dermatobranchus striatellus* Baba,1949 | Tropical–subtropical species |
|  | Nudibranchia | Cladobranchia |  | Bornellidae |  | Bornella | *Bornella stellifer*  (A.Adams&Reeve in A.Adams,1848) | Tropical–subtropical species |
|  | Nudibranchia | Cladobranchia |  | Dotidae |  | Doto | *Doto japonica* Odhner,1936 | Temperate species |
|  | Nudibranchia | Cladobranchia |  | Dotidae |  | Doto | *Doto bella* Baba,1938 | Temperate species |
|  | Nudibranchia | Cladobranchia | Fionidea | Fionidae |  | Eubramchus | *Eubranchus virginalis*  (Baba,1949) | Tropical–subtropical species |
|  | Nudibranchia | Cladobranchia | Fionidea | Fionidae |  | Eubramchus | *Eubranchus inabai* Baba,1964 | Tropical–subtropical species |
|  | Nudibranchia | Cladobranchia | Fionidea | Fionidae |  | Eubramchus | *Eubranchus horii* Baba,1960 | Temperate species |
|  | Nudibranchia | Cladobranchia | Fionidea | Fionidae |  | Leostyletus | *Leostyletus misakiensis* (Baba,1960) | Subarctic–arctic species |
|  | Nudibranchia | Cladobranchia | Fionidea | Fionidae |  | Tenellia | *Tenellia futairo*(Baba,1963) | Temperate species |
|  | Nudibranchia | Cladobranchia | Fionidea | Trinchesiidae |  | Catriona | *Catriona pinnifera* (Baba, 1949) | Tropical–subtropical species |
|  | Nudibranchia | Cladobranchia | Fionidea | Fionidae |  | Tenellia | *Tenellia ornata*(Baba,1937) | Tropical–subtropical species |
|  | Nudibranchia | Cladobranchia | Fionidea | Fionidae |  | Tenellia | *Tenellia pupillae*(Baba,1961) | Temperate species |
|  | Nudibranchia |  | Aeolidioidea | Facelinidae |  | Cratena | *Cratena lineata*(Eliot,1904) | Tropical–subtropical species |
|  | Nudibranchia |  | Aeolidioidea | Facelinidae |  | Herviella | *Herviella affinis* Baba,1960 | Temperate species |
|  | Nudibranchia |  | Aeolidioidea | Facelinidae |  | Herviella | *Herviella yatsui*(Baba,1930) | Tropical–subtropical species |
|  | Nudibranchia |  | Aeolidioidea | Facelinidae |  | Sakuraeolis | *Sakuraeolis enosimensis*  (Baba,1930) | Temperate species |
|  | Nudibranchia |  | Aeolidioidea | Aeolidiidae |  | Anteaeolidiella | *Anteaeolidiella takanosimensis*  (Baba,1930) | Temperate species |
|  | Nudibranchia |  | Aeolidioidea | Aeolidiidae |  | Limenandra | *Limenandra fusiformis*  (Baba,1949) | Tropical–subtropical species |
|  | Nudibranchia |  | Aeolidioidea | Aeolidiidae |  | Bulbaeolidia | *Bullbaeolidia japonica*  (Eliot,1913) | Temperate species |
| Takahama Ingeri-bana Cape | Sacoglossa |  |  | Limapontiidae |  | Ercolania | *Ercolania boodleae*(Baba,1938) | Subarctic–arctic species |
|  | Sacoglossa |  |  | Hermaeidae |  | Aplysiopsis | *Aplysiopsis nigra*(Baba,1949) | Temperate species |
|  | Sacoglossa |  |  | Limapontiidae |  | Placida | *Placida* sp.1 | Temperate species |
|  | Sacoglossa |  |  | Plakobranchidae |  | Elysia | *Elysia atroviridis* Baba,1955 | Temperate species |
|  | Sacoglossa |  |  | Plakobranchidae |  | Elysia | *Elysia amakusana* Baba,1955 | Tropical–subtropical species |
| Igeshuku,MeotoIwa | Sacoglossa |  |  | Limapontiidae |  | Placida | *Placida kevinleei* McCarthy,Krug&Valdes,2017 | Tropical–subtropical–temperate species |
|  | Sacoglossa |  |  | Oxynoidae |  | Oxynoe | *Oxynoe viridis*(Pease,1861) | Tropical–subtropical–temperate species |
|  | Sacoglossa |  |  | Plakobranchidae |  | Elysia | *Elysia obtusa* Baba,1938 | Tropical–subtropical–temperate species |

(b) From Kawahara (unpublished).

| **Site** | **date** | **Order** | **Suborder** | **Superfamily** | **Family** | **Subfamily** | **Genus** | **Species** | **climate classification** | **Number of individuals** |
| --- | --- | --- | --- | --- | --- | --- | --- | --- | --- | --- |
| Tatsunokuchi | 2001-05-04 | Nudibranchia | Doridacea(Cryptobranchia) |  | Chromodorididae |  | Doriprismatica | *Doriprismatica atromarginata*  (Cuvier,1804) | Tropical–subtropical–temperate species | 1 |
| Tatsunokuchi |  | Nudibranchia | Doridacea(Cryptobranchia) |  | Chromodorididae |  | Hypselodoris | *Hypselodoris festiva*  (A.Adams,1861) | Temperate species | 1 |
| Tatsunokuchi |  | Nudibranchia | Doridacea(Cryptobranchia) |  | Chromodorididae |  | Verconia | *Verconia subnivalis*(Baba,1987) | Temperate species | 1 |
| Tatsunokuchi |  | Nudibranchia | Doridacea(Cryptobranchia) |  | Discodorididae |  | Jorunna | *Jorunna parva*(Baba,1938) | Tropical–subtropical species | 1 |
| Tatsunokuchi |  | Nudibranchia | Doridacea(Porostomata) |  | Dendorodorididae |  | Dendrodoris | *Dendrodoris denisoni*  (Angas,1864) | Tropical–subtropical–temperate species | 1 |
| Tatsunokuchi |  | Nudibranchia | Cladobranchia |  | Aminidae |  | Dematobranchus | *Dermatobranchus primus* Baba,1976 | Temperate species | 1 |
| Tatsunokuchi |  | Nudibranchia |  | Fionidea | Fionidae |  | Tenellia | *Tenellia diversicolor*(Baba,1975) | Tropical–subtropical species | 1 |
| Tatsunokuchi | 2001-06-24 | Sacoglossa |  |  | Plakobranchidae |  | Elysia | *Elysia atroviridis* Baba,1955 | Temperate species | 2 |
| Tatsunokuchi |  | Nudibranchia | Doridacea(Cryptobranchia) |  | Chromodorididae |  | Goniobranchus | *Goniobranchus orientails*  (Rudman,1983) | Tropical–subtropical species | 1 |
| Tatsunokuchi | 2001-06-30 | Nudibranchia | Doridacea(Cryptobranchia) |  | Chromodorididae |  | Goniobranchus | *Goniobranchus orientails*  (Rudman,1983) | Tropical–subtropical species | 2 |
| Tatsunokuchi |  | Nudibranchia | Doridacea(Cryptobranchia) |  | Chromodorididae |  | Verconia | *Verconia purpurea*(Baba,1949) | Temperate species | 1 |
| Tatsunokuchi |  | Nudibranchia | Doridacea(Porostomata) |  | Phyllidiidae |  | Phyllidiella | *Phyllidiella pustulosa*  (Cuvier,1804) | Tropical–subtropical–temperate species | 1 |
| Tatsunokuchi |  | Nudibranchia | Doridacea(Cryptobranchia) |  | Chromodorididae |  | Verconia | *Verconia norba*  (Er.Marcus&Ev.Marcus,1970) | Tropical–subtropical–temperate species | 1 |
| Tatsunokuchi | 2001-07-05 | Nudibranchia | Doridacea(Cryptobranchia) |  | Chromodorididae |  | Goniobranchus | *Goniobranchus orientails*  (Rudman,1983) | Tropical–subtropical species | 3 |
| Tatsunokuchi |  | Nudibranchia | Doridacea(Cryptobranchia) |  | Chromodorididae |  | Doriprismatica | *Doriprismatica atromarginata*  (Cuvier,1804) | Tropical–subtropical–temperate species | 4 |
| Tatsunokuchi |  | Nudibranchia | Doridacea(Cryptobranchia) |  | Chromodorididae |  | Hypselodoris | *Hypselodoris sagamiensis*  (Baba,1949) | Tropical–subtropical species | 3 |
| Tatsunokuchi |  | Nudibranchia | Doridacea(Cryptobranchia) |  | Chromodorididae |  | Verconia | *Verconia purpurea*(Baba,1949) | Temperate species | 4 |
| Tatsunokuchi |  | Nudibranchia | Doridacea(Porostomata) |  | Phyllidiidae |  | Phyllidiella | *Phyllidiella pustulosa*  (Cuvier,1804) | Tropical–subtropical–temperate species | 3 |
| Tatsunokuchi | 2001-07-20 | Nudibranchia | Doridacea(Phanerobranchia) |  | Gymnodoridae |  | Gymnodoris | *Gymnodoris impudica*  (Ruppell&Leuckart,1828) | Tropical–subtropical species | 1 |
| Tatsunokuchi |  | Nudibranchia | Doridacea(Cryptobranchia) |  | Chromodorididae |  | Doriprismatica | *Doriprismatica atromarginata*  (Cuvier,1804) | Tropical–subtropical–temperate species | 3 |
| Tatsunokuchi |  | Nudibranchia | Doridacea(Cryptobranchia) |  | Chromodorididae |  | Hypselodoris | *Hypselodoris festiva*  (A.Adams,1861) | Temperate species | 1 |
| Tatsunokuchi | 2001-08-02 | Nudibranchia | Doridacea(Cryptobranchia) |  | Chromodorididae |  | Doriprismatica | *Doriprismatica atromarginata*  (Cuvier,1804) | Tropical–subtropical–temperate species | 1 |
| Tatsunokuchi | 2001-10-02 |  |  |  |  |  |  | None observed |  |  |
| Tatsunokuchi | 2001-12-15 |  |  |  |  |  |  | None observed |  |  |
| Tatsunokuchi | 2002-04-26 | Aplysiida |  |  | Aplysiidae |  | Aplysia | *Aplysia kurodai*(Baba,1937) | Tropical–subtropical species | 12 |
| Tatsunokuchi | 2002-04-29 | Aplysiida |  |  | Aplysiidae |  | Aplysia | *Aplysia kurodai*(Baba,1937) | Tropical–subtropical species | 117 |
| Tatsunokuchi |  | Aplysiida |  |  | Aplysiidae |  | Aplysia | *Aplysia japonica*  (G.B.Sowerby Ⅱ,1869) | Temperate species | 1 |
| Tatsunokuchi |  | Nudibranchia | Doridacea(Phanerobranchia) |  | Goniodorididae |  | Okenia | *Okenia hiroi*(Baba,1938) | Tropical–subtropical species | 1 |
| Tatsunokuchi | 2002-05-13 | Aplysiida |  |  | Aplysiidae |  | Aplysia | *Aplysia kurodai*(Baba,1937) | Tropical–subtropical species | 50 |
| Tatsunokuchi |  | Sacoglossa |  |  | Limapontiidae |  | Placida | *Placida* sp.1 | Temperate species | 1 |
| Tatsunokuchi |  | Sacoglossa |  |  | Plakobranchidae |  | Elysia | *Elysia trisinuata* Baba,1949 | Tropical–subtropical species | 1 |
| Tatsunokuchi |  | Nudibranchia | Doridacea(Cryptobranchia) |  | Chromodorididae |  | Goniobranchus | *Goniobranchus orientails*  (Rudman,1983) | Tropical–subtropical species | 1 |
| Tatsunokuchi |  | Nudibranchia | Doridacea(Cryptobranchia) |  | Chromodorididae |  | Goniobranchus | *Goniobranchus tinctorius*  (Ruppell&Leuckart,1830) | Tropical–subtropical species | 1 |
| Tatsunokuchi |  | Nudibranchia | Doridacea(Cryptobranchia) |  | Chromodorididae |  | Hypselodoris | *Hypselodoris festiva*  (A.Adams,1861) | Temperate species | 1 |
| Tatsunokuchi | 2002-06-05 | Nudibranchia | Doridacea(Phanerobranchia) |  | Goniodorididae |  | Okenia | *Okenia hiroi*(Baba,1938) | Tropical–subtropical species | 5 |
| Tatsunokuchi |  | Nudibranchia | Doridacea(Porostomata) |  | Dendorodorididae |  | Dendrodoris | *Dendrodoris denisoni*  (Angas,1864) | Tropical–subtropical–temperate species | 1 |
| Tatsunokuchi | 2002-06-28 | Sacoglossa |  |  | Plakobranchidae |  | Elysia | *Elysia trisinuata* Baba,1949 | Tropical–subtropical species | 1 |
| Tatsunokuchi |  | Nudibranchia | Doridacea(Cryptobranchia) |  | Chromodorididae |  | Hypselodoris | *Hypselodoris festiva*  (A.Adams,1861) | Temperate species | 1 |
| Tatsunokuchi | 2002-07-12 | Sacoglossa |  |  | Plakobranchidae |  | Elysia | *Elysia trisinuata* Baba,1949 | Tropical–subtropical species | 2 |
| Tatsunokuchi | 2002-08-06 |  |  |  |  |  |  | None observed |  |  |
| Tatsunokuchi | 2002-08-15 |  |  |  |  |  |  | None observed |  |  |
| Tatsunokuchi | 2002-09-18 | Sacoglossa |  |  | Elysiidae |  | Elysia | *Elysia marginata*(Pease,1871) | Tropical–subtropical–temperate species | 1 |
| Tatsunokuchi | 2002-09-24 | Nudibranchia | Doridacea(Cryptobranchia) |  | Chromodorididae |  | Doriprismatica | *Doriprismatica atromarginata*  (Cuvier,1804) | Tropical–subtropical–temperate species | 1 |
| Tatsunokuchi |  | Nudibranchia | Doridacea(Porostomata) |  | Dendorodorididae |  | Dendrodoris | *Dendrodoris denisoni*  (Angas,1864) | Tropical–subtropical–temperate species | 1 |
| Tatsunokuchi | 2002-10-09 |  |  |  |  |  |  | None observed |  |  |
| Tatsunokuchi | 2002-10-17 |  |  |  |  |  |  | None observed |  |  |
| Tatsunokuchi | 2002-11-28 | Sacoglossa |  |  | Elysiidae |  | Elysia | *Elysia marginata*(Pease,1871) | Tropical–subtropical–temperate species | 1 |
| Tatsunokuchi |  | Nudibranchia | Doridacea(Phanerobranchia) |  | Polyceridae | Polycerinae | Polycera | *Polycera japonica* Baba,1949 | Tropical–subtropical–temperate species | 1 |
| Tatsunokuchi | 2002-12-06 | Sacoglossa |  |  | Limapontiidae |  | Placida | *Placida kevinleei*  McCarthy,Krug&Valdes,2017 | Tropical–subtropical–temperate species | 9 |
| Tatsunokuchi |  | Nudibranchia | Doridacea(Porostomata) |  | Phyllidiidae |  | Phyllidiella | *Phyllidiella pustulosa*  (Cuvier,1804) | Tropical–subtropical–temperate species | 1 |
| Tatsunokuchi | 2003-03-29 | Sacoglossa |  |  | Limapontiidae |  | Placida | *Placida* sp.1 | Temperate species | 1 |
| Tatsunokuchi | 2003-03-30 | Aplysiida |  |  | Aplysiidae |  | Aplysia | *Aplysia kurodai*(Baba,1937) | Tropical–subtropical species | 1 |
| Tatsunokuchi |  | Sacoglossa |  |  | Limapontiidae |  | Placida | *Placida* sp.1 | Temperate species | 14 |
| Tatsunokuchi |  | Sacoglossa |  |  | Hermaeidae |  | Aplysiopsis | *Aplysiopsis minor*(Baba,1959) | Temperate species | 4 |
| Tatsunokuchi |  | Sacoglossa |  |  | Limapontiidae |  | Ercolania | *Ercolania boodleae*(Baba,1938) | Tropical–subtropical species | 24 |
| Tatsunokuchi |  | Sacoglossa |  |  | Plakobranchidae |  | Elysia | *Elysia atroviridis* Baba,1955 | Temperate species | 3 |
| Tatsunokuchi | 2003-05-21 | Aplysiida |  |  | Aplysiidae |  | Aplysia | *Aplysia kurodai*(Baba,1937) | Tropical–subtropical species | 2 |
| Tatsunokuchi | 2003-06-03 | Nudibranchia | Doridacea(Cryptobranchia) |  | Chromodorididae |  | Goniobranchus | *Goniobranchus orientails*  (Rudman,1983) | Tropical–subtropical species | 1 |
| Tatsunokuchi | 2003-07-23 | Nudibranchia | Doridacea(Cryptobranchia) |  | Chromodorididae |  | Doriprismatica | *Doriprismatica atromarginata*  (Cuvier,1804) | Tropical–subtropical–temperate species | 1 |
| Tatsunokuchi | 2003-09-24 | Sacoglossa |  |  | Elysiidae |  | Elysia | *Elysia marginata*(Pease,1871) | Tropical–subtropical–temperate species | 8 |
| Tatsunokuchi |  | Nudibranchia | Doridacea(Cryptobranchia) |  | Chromodorididae |  | Doriprismatica | *Doriprismatica atromarginata*  (Cuvier,1804) | Tropical–subtropical–temperate species | 3 |
| Nomozaki-Akase | 2001-05-05 | Nudibranchia | Doridacea(Phanerobranchia) |  | Goniodorididae |  | Okenia | *Okenia hiroi*(Baba,1938) | Tropical–subtropical species | 1 |
| Nomozaki-Akase |  | Nudibranchia | Doridacea(Cryptobranchia) |  | Chromodorididae |  | Goniobranchus | *Goniobranchus orientails*  (Rudman,1983) | Tropical–subtropical species | 3 |
| Nomozaki-Akase |  | Nudibranchia | Doridacea(Cryptobranchia) |  | Chromodorididae |  | Goniobranchus | *Goniobranchus tinctorius*  (Ruppell&Leuckart,1830) | Tropical–subtropical species | 1 |
| Nomozaki-Akase |  | Nudibranchia | Doridacea(Cryptobranchia) |  | Chromodorididae |  | Hypselodoris | *Hypselodoris festiva*  (A.Adams,1861) | Temperate species | 6 |
| Nomozaki-Akase |  | Nudibranchia | Doridacea(Porostomata) |  | Dendorodorididae |  | Dendrodoris | *Dendrodoris denisoni*  (Angas,1864) | Tropical–subtropical–temperate species | 1 |
| Nomozaki-Akase | 2001-06-09 | Nudibranchia | Doridacea(Cryptobranchia) |  | Chromodorididae |  | Goniobranchus | *Goniobranchus orientails*  (Rudman,1983) | Tropical–subtropical species | 1 |
| Nomozaki-Akase |  | Nudibranchia | Doridacea(Porostomata) |  | Dendorodorididae |  | Dendrodoris | *Dendrodoris denisoni*  (Angas,1864) | Tropical–subtropical–temperate species | 3 |
| Nomozaki-Akase |  | Nudibranchia | Cladobranchia |  | Arminidae |  | Dermatobranchus | *Dermatobranchus striatellus*  Baba,1949 | Tropical–subtropical species | 3 |
| Nomozaki-Akase | 2001-07-31 | Nudibranchia | Doridacea(Cryptobranchia) |  | Chromodorididae |  | Hypselodoris | *Hypselodoris festiva*  (A.Adams,1861) | Temperate species | 1 |
| Nomozaki-Akase |  | Nudibranchia | Doridacea(Cryptobranchia) |  | Chromodorididae |  | Doriprismatica | *Doriprismatica atromarginata*  (Cuvier,1804) | Tropical–subtropical–temperate species | 1 |
| Nomozaki-Akase |  | Nudibranchia | Doridacea(Porostomata) |  | Phyllidiidae |  | Phyllidiella | *Phyllidiella pustulosa*  (Cuvier,1804) | Tropical–subtropical–temperate species | 1 |
| Nomozaki-Akase |  | Nudibranchia | Cladobranchia |  | Tritoniidae |  | Tritoniopsis | *Tritoniopsis elegans*  (Audouin in Savigny,1826) | Tropical–subtropical species | 1 |
| Nomozaki-Akase | 2001-08-04 | Aplysiida |  |  | Aplysiidae |  | Aplysia | *Aplysia kurodai(*Baba,1937) | Tropical–subtropical species | 4 |
| Nomozaki-Akase |  | Nudibranchia | Doridacea(Cryptobranchia) |  | Chromodorididae |  | Goniobranchus | *Goniobranchus orientails*  (Rudman,1983) | Tropical–subtropical species | 4 |
| Nomozaki-Akase |  | Nudibranchia | Doridacea(Cryptobranchia) |  | Chromodorididae |  | Goniobranchus | *Goniobranchus fidelis*  (Kelaart,1858) | Tropical–subtropical–temperate species | 2 |
| Nomozaki-Akase |  | Nudibranchia | Doridacea(Cryptobranchia) |  | Chromodorididae |  | Goniobranchus | *Goniobranchus sinensis*  (Rudman,1985) | Tropical–subtropical species | 1 |
| Nomozaki-Akase |  | Nudibranchia | Doridacea(Porostomata) |  | Dendorodorididae |  | Dendrodoris | *Dendrodoris guttata*  (Odhner,1917) | Tropical–subtropical species | 2 |
| Nomozaki-Akase | 2001-08-08 | Nudibranchia | Doridacea(Cryptobranchia) |  | Actinocyclidae |  | Actynocyclus | *Actinocyclus papillatus*  (Bergh,1878) | Tropical–subtropical species | 1 |
| Nomozaki-Akase | 2001-08-31 | Nudibranchia |  | Fionidea | Samlidae |  | Samla | *Samla takashigei*  Korshunova et al.,2017 | Tropical–subtropical species | 5 |
| Nomozaki-Akase | 2001-09-02 | Nudibranchia | Doridacea(Porostomata) |  | Dendorodorididae |  | Dendrodoris | *Dendrodoris denisoni*  (Angas,1864) | Tropical–subtropical–temperate species | 3 |
| Nomozaki-Akase | 2001-09-04 | Nudibranchia |  | Fionidea | Samlidae |  | Samla | *Samla takashigei*  Korshunova et al.,2017 | Tropical–subtropical species | 3 |
| Nomozaki-Akase | 2001-10-10 |  |  |  |  |  |  | None observed |  |  |
| Nomozaki-Akase | 2002-04-03 | Cephalaspidea |  |  | Haminoeidae |  | Atys | *Atys semistriatus*  Pease,1860 | Tropical–subtropical–temperate species | 2 |
| Nomozaki-Akase |  | Aplysiida |  |  | Aplysiidae |  | Aplysia | *Aplysia oculifera*  (Adams&Reeve,1850) | Tropical–subtropical species | 12 |
| Nomozaki-Akase |  | Aplysiida |  |  | Aplysiidae |  | Aplysia | *Aplysia japonica*  (G.B.Sowerby Ⅱ,1869) | Temperate species | 1 |
| Nomozaki-Akase |  | Nudibranchia |  | Aeolidioidea | Facelinidae |  | Sakuraeolis | *Sakuraeolis enosimensis*  (Baba,1930) | Temperate species | 1 |
| Nomozaki-Akase | 2002-05-02 | Aplysiida |  |  | Aplysiidae |  | Aplysia | *Aplysia japonica*  (G.B.Sowerby Ⅱ,1869) | Temperate species | 2 |
| Nomozaki-Akase | 2002-05-04 | Aplysiida |  |  | Aplysiidae |  | Aplysia | *Aplysia kurodai*(Baba,1937) | Tropical–subtropical species | 50 |
| Nomozaki-Akase |  | Aplysiida |  |  | Aplysiidae |  | Aplysia | *Aplysia oculifera*  (Adams&Reeve,1850) | Tropical–subtropical species | 50 |
| Nomozaki-Akase |  | Nudibranchia | Doridacea(Cryptobranchia) |  | Chromodorididae |  | Goniobranchus | *Goniobranchus orientails*  (Rudman,1983) | Tropical–subtropical species | 2 |
| Nomozaki-Akase |  | Nudibranchia | Doridacea(Cryptobranchia) |  | Discodorididae |  | Rostanga | *Rostanga orientalis*  Rudman&Avern,1989 | Temperate species | 1 |
| Nomozaki-Akase |  | Nudibranchia | Doridacea(Cryptobranchia) |  | Discodorididae |  | Jorunna | *Jorunna parva*(Baba,1938) | Tropical–subtropical species | 1 |
| Nomozaki-Akase | 2002-06-01 | Nudibranchia | Doridacea(Cryptobranchia) |  | Chromodorididae |  | Goniobranchus | *Goniobranchus orientails*  (Rudman,1983) | Tropical–subtropical species | 1 |
| Nomozaki-Akase | 2002-08-27 | Sacoglossa |  |  | Elysiidae |  | Elysia | *Elysia marginata*  (Pease,1871) | Tropical–subtropical–temperate species | 14 |
| Nomozaki-Akase |  | Nudibranchia | Doridacea(Porostomata) |  | Dendorodorididae |  | Dendrodoris | *Dendrodoris denisoni*  (Angas,1864) | Tropical–subtropical–temperate species | 1 |
| Nomozaki-Akase |  | Nudibranchia |  | Fionidea | Samlidae |  | Samla | *Samla takashigei*  Korshunova et al.,2017 | Tropical–subtropical species | 5 |
| Nomozaki-Akase | 2002-10-24 |  |  |  |  |  |  | None observed |  |  |

**Table S2** Summary of environmental data recorded during the present scuba diving surveys.

| Survey Date | Site | Method | Start | End | Duration  （min） | Air  (℃) | Water  (℃) | Mean Depth  （ｍ） | Max Depth  （ｍ） | Visibility  (m) | Tide | Sea Conditions | Weather |
| --- | --- | --- | --- | --- | --- | --- | --- | --- | --- | --- | --- | --- | --- |
| 2023-06-02 | Tatsunokuchi | Scuba diving | 11:39 | 12:46 | 67 | 18 | 20.8 | 3.6 | 5.5 | 5 | Medium tide | Calm | Rainy |
| 2023-07-05 | Nomozaki-Akase | Scuba diving | 9:54 | 11:15 | 81 | 25 | 23.8 | 4.0 | 7.2 | 8 | Medium tide  (at low tide) | Calm | Rainy |
| 2023-07-05 | Nomozaki-Akase | Scuba diving | 11:51 | 12:50 | 51 | 27 | 23.8 | 2.9 | 5.3 | 8 | Medium tide  (at low tide) | Calm | Rainy |
| 2023-07-19 | Nomozaki-Akase | Scuba diving | 9:19 | 10:40 | 81 | 26 | 25.9 | 3.6 | 5.3 | 8 | Medium tide  (at low tide) | Slightly wavy | Rainy |
| 2023-07-19 | Nomozaki-Akase | Scuba diving | 11:11 | 12:22 | 72 | 27 | 25.9 | 4.1 | 7.4 | 8 | Spring tide  (ebb tide) | Slightly wavy | Rainy |
| 2023-07-31 | Tatsunokuchi | Scuba diving | 9:01 | 9:56 | 55 | 35 | 28.6 | 4.2 | 6.2 | 5 | Neap tide | Calm | Rainy |
| 2023-07-31 | Tatsunokuchi | Scuba diving | 12:06 | 13:02 | 56 | 35 | 27.2 | 6.9 | 14.3 | 5 | Neap tide | Calm | Rainy |
| 2023-08-02 | Nomozaki-Akase | Scuba diving | 9:28 | 10:34 | 66 | 33 | 26.5 | 3.9 | 7.9 | 8 | Spring tide  (ebb tide) | Wavy | Sunny |
| 2023-08-02 | Nomozaki-Akase | Scuba diving | 11:07 | 12:09 | 62 | 33 | 27.2 | 2.3 | 3.8 | 8 | Spring tide  (ebb tide) | Wavy | Sunny |
| 2023-08-29 | Tatsunokuchi | Scuba diving | 8:59 | 9:56 | 59 | 31 | 25.8 | 7.1 | 13.6 | 3 | Spring tide  (ebb tide) | Calm | Sunny |
| 2023-08-29 | Tatsunokuchi | Scuba diving | 10:53 | 11:48 | 55 | 31 | 27.6 | 3.5 | 5.5 | 3 | Spring tide (  ebb tide) | Calm | Cloudy |
| 2023-09-19 | Tatsunokuchi | Scuba diving | 9:37 | 10:38 | 61 | 26 | 27.7 | 7.1 | 10.8 | 5 | Spring tide | Calm | Cloudy |
| 2023-09-19 | Tatsunokuchi | Scuba diving | 11:34 | 12:40 | 66 | 26 | 27.4 | 4.8 | 8.6 | 5 | Spring tide | Calm | Sunny |
| 2023-09-29 | Nomozaki-Akase | Scuba diving | 9:49 | 11:30 | 73 | 25 | 26.9 | 3.5 | 5.7 | 8 | Spring tide | Calm | Sunny |
| 2023-09-29 | Nomozaki-Akase | Scuba diving | 11:43 | 12:31 | 48 | 28 | 27 | 2.7 | 5.2 | 8 | Spring tide | Calm | Sunny |
| 2023-10-03 | Nomozaki-Akase | Scuba diving | 9:30 | 10:26 | 56 | 23 | 25.9 | 3.7 | 5.3 | 8 | Medium tide  (ebb tide) | Calm | Sunny |
| 2023-10-03 | Nomozaki-Akase | Scuba diving | 11:10 | 12:25 | 75 | 25 | 25.8 | 4.4 | 7.4 | 8 | Medium tide  (ebb tide) | Calm | Sunny |
| 2023-10-23 | Tatsunokuchi | Scuba diving | 9:41 | 10:35 | 54 | 20 | 23.5 | 9.6 | 14.3 | 5 | Neap tide | Calm | Sunny |
| 2023-10-23 | Tatsunokuchi | Scuba diving | 11:26 | 12:06 | 40 | 21 | 22.9 | 4.1 | 6.7 | 5 | Neap tide | Calm | Sunny |
| 2023-11-16 | Nomozaki-Akase | Scuba diving | 9:38 | 10:31 | 53 | 13 | 20.4 | 3.3 | 5.5 | 8 | Medium tide  (at high tide) | Calm | Cloudy |
| 2023-11-24 | Tatsunokuchi | Scuba diving | 9:19 | 10:03 | 44 | 13 | 18.4 | 4.5 | 5.7 | 4 | Medium tide  (at low tide) | Calm | Cloudy |
| 2023-01-16 | Nomozaki-Akase | Scuba diving | 9:57 | 10:53 | 56 | 7 | 14.5 | 3.9 | 5.8 | 8 | Medium tide  (at high tide) | Wavy | Sunny |
| 2023-01-16 | Nomozaki-Akase | Scuba diving | 11:45 | 12:42 | 57 | 9 | 15.2 | 3.3 | 4.2 | 8 | Medium tide  (at high tide) | Wavy | Sunny |
| 2024-01-19 | Tatsunokuchi | Scuba diving | 9:19 | 10:26 | 67 | 12 | 15.4 | 12.4 | 19.4 | 5 | Neap tide | Calm | Rainy |
| 2024-01-19 | Tatsunokuchi | Scuba diving | 11:36 | 12:48 | 72 | 14 | 15.2 | 4.9 | 7.7 | 5 | Neap tide | Calm | Cloudy |

**Table S3** Summary of nudibranch assemblages recorded during the present scuba diving surveys.

| **Site** | **date** | **Order** | **Suborder** | **Superfamily** | **Family** | **Subfamily** | **Genus** | **Species** | **climate classification** | **Number of individuals** |
| --- | --- | --- | --- | --- | --- | --- | --- | --- | --- | --- |
| Nomozaki-Akase | 2023-07-05 | Aplysiida |  |  | Aplysiidae |  | Aplysia | *Aplysia japonica*  (G.B.Sowerby Ⅱ,1869) | Temperate species | 3 |
| Nomozaki-Akase |  | Sacoglossa |  |  | Plakobranchidae |  | Elysia | *Elysia trisinuata* Baba,1949 | Tropical–subtropical species | 3 |
| Nomozaki-Akase |  | Nudibranchia | Doridacea(Cryptobranchia) |  | Chromodorididae |  | Doriprismatica | *Doriprismatica atromarginata*  (Cuvier,1804) | Tropical–subtropical–temperate species | 2 |
| Nomozaki-Akase |  | Nudibranchia | Doridacea(Cryptobranchia) |  | Chromodorididae |  | Goniobranchus | *Goniobranchus orientails*  (Rudman,1983) | Tropical–subtropical species | 2 |
| Nomozaki-Akase |  | Nudibranchia | Doridacea(Cryptobranchia) |  | Chromodorididae |  | Goniobranchus | *Goniobranchus sinensis*  (Rudman,1985) | Tropical–subtropical species | 1 |
| Nomozaki-Akase |  | Nudibranchia | Doridacea(Cryptobranchia) |  | Chromodorididae |  | Goniobranchus | *Goniobranchus tinctorius*  (Ruppell&Leuckart,1830) | Tropical–subtropical species | 2 |
| Nomozaki-Akase |  | Nudibranchia | Doridacea(Cryptobranchia) |  | Chromodorididae |  | Hypselodoris | *Hypselodoris festiva*  (A.Adams,1861) | Temperate species | 2 |
| Nomozaki-Akase |  |  | Doridacea(Cryptobranchia) |  | Chromodorididae |  | Hypselodoris | *Hypselodoris placida*(Baba,1949) | Temperate species | 1 |
| Nomozaki-Akase |  | Nudibranchia | Doridacea(Porostomata) |  | Dendorodorididae |  | Dendrodoris | *Dendrodoris denisoni*  (Angas,1864) | Tropical–subtropical–temperate species | 1 |
| Nomozaki-Akase |  | Nudibranchia | Cladobranchia |  | Tritoniidae |  | Tritoniopsis | *Tritoniopsis elegans*  (Audouin in Savigny,1826) | Tropical–subtropical species | 2 |
| Nomozaki-Akase | 2023-07-19 | Aplysiida |  |  | Aplysiidae |  | Aplysia | *Aplysia japonica*  (G.B.Sowerby Ⅱ,1869) | Temperate species | 3 |
| Nomozaki-Akase |  | Sacoglossa |  |  | Plakobranchidae |  | Elysia | *Elysia trisinuata* Baba,1949 | Tropical–subtropical species | 1 |
| Nomozaki-Akase |  | Sacoglossa |  |  | Plakobranchidae |  | Thuridilla | *Thuridilla albopustulosa* Gosliner,1995 | Tropical–subtropical species | 1 |
| Nomozaki-Akase |  | Sacoglossa |  |  | Plakobranchidae |  | Thuridilla | *Thuridilla splendens*(Baba,1949) | Tropical–subtropical species | 1 |
| Nomozaki-Akase |  | Nudibranchia | Doridacea(Cryptobranchia) |  | Chromodorididae |  | Doriprismatica | *Doriprismatica atromarginata*  (Cuvier,1804) | Tropical–subtropical–temperate species | 1 |
| Nomozaki-Akase |  | Nudibranchia | Doridacea(Cryptobranchia) |  | Chromodorididae |  | Goniobranchus | *Goniobranchus fidelis*  (Kelaart,1858) | Tropical–subtropical–temperate species | 1 |
| Nomozaki-Akase |  | Nudibranchia | Doridacea(Cryptobranchia) |  | Chromodorididae |  | Goniobranchus | *Goniobranchus orientails*  (Rudman,1983) | Tropical–subtropical species | 5 |
| Nomozaki-Akase |  | Nudibranchia | Doridacea(Cryptobranchia) |  | Chromodorididae |  | Goniobranchus | *Goniobranchus sinensis*  (Rudman,1985) | Tropical–subtropical species | 3 |
| Nomozaki-Akase |  | Nudibranchia | Doridacea(Cryptobranchia) |  | Chromodorididae |  | Hypselodoris | *Hypselodoris festiva*  (A.Adams,1861) | Temperate species | 3 |
| Nomozaki-Akase |  | Nudibranchia | Doridacea(Cryptobranchia) |  | Chromodorididae |  | Hypselodoris | *Hypselodoris sagamiensis*  (Baba,1949) | Tropical–subtropical species | 1 |
| Nomozaki-Akase |  | Nudibranchia | Cladobranchia |  | Tritoniidae |  | Tritoniopsis | *Tritoniopsis elegans*  (Audouin in Savigny,1826) | Tropical–subtropical species | 1 |
| Nomozaki-Akase | 2023-08-02 | Aplysiida |  |  | Aplysiidae |  | Aplysia | *Aplysia japonica*  (G.B.Sowerby Ⅱ,1869) | Temperate species | 2 |
| Nomozaki-Akase |  | Aplysiida |  |  | Aplysiidae |  | Aplysia | *Aplysia kurodai*(Baba,1937) | Tropical–subtropical species | 3 |
| Nomozaki-Akase |  | Sacoglossa |  |  | Plakobranchidae |  | Elysia | *Elysia trisinuata* Baba,1949 | Tropical–subtropical species | 1 |
| Nomozaki-Akase |  | Nudibranchia | Doridacea(Cryptobranchia) |  | Chromodorididae |  | Doriprismatica | *Doriprismatica atromarginata*  (Cuvier,1804) | Tropical–subtropical–temperate species | 2 |
| Nomozaki-Akase |  | Nudibranchia | Doridacea(Cryptobranchia) |  | Chromodorididae |  | Goniobranchus | *Goniobranchus aureopurpureus*  (Collingwood,1881) | Tropical–subtropical species | 2 |
| Nomozaki-Akase |  | Nudibranchia | Doridacea(Cryptobranchia) |  | Chromodorididae |  | Verconia | *Verconia nivalis*(Baba,1937) | Temperate species | 1 |
| Nomozaki-Akase |  | Nudibranchia | Doridacea(Cryptobranchia) |  | Chromodorididae |  | Goniobranchus | *Goniobranchus sinensis*  (Rudman,1985) | Tropical–subtropical species | 1 |
| Nomozaki-Akase |  | Nudibranchia | Doridacea(Cryptobranchia) |  | Chromodorididae |  | Hypselodoris | *Hypselodoris festiva*  (A.Adams,1861) | Temperate species | 1 |
| Nomozaki-Akase |  | Nudibranchia | Doridacea(Porostomata) |  | Dendorodorididae |  | Dendrodoris | *Dendrodoris denisoni*  (Angas,1864) | Tropical–subtropical–temperate species | 1 |
| Nomozaki-Akase |  | Nudibranchia | Cladobranchia |  | Tritoniidae |  | Tritoniopsis | *Tritoniopsis elegans*  (Audouin in Savigny,1826) | Tropical–subtropical species | 3 |
| Nomozaki-Akase | 2023-09-29 | Nudibranchia | Doridacea(Cryptobranchia) |  | Chromodorididae |  | Goniobranchus | *Goniobranchus tinctorius*  (Ruppell&Leuckart,1830) | Tropical–subtropical species | 1 |
| Nomozaki-Akase |  | Nudibranchia | Cladobranchia |  | Tritoniidae |  | Tritoniopsis | *Tritoniopsis elegans*  (Audouin in Savigny,1826) | Tropical–subtropical species | 1 |
| Nomozaki-Akase |  | Nudibranchia | Cladobranchia | Aeolidioidea | Facelinidae |  | Phyllodesmium | *Phyllodesmium magnum* Rudman,1991 | Tropical–subtropical–temperate species | 1 |
| Nomozaki-Akase | 2023-10-03 | Sacoglossa |  |  | Plakobranchidae |  | Thuridilla | *Thuridilla splendens*(Baba,1949) | Tropical–subtropical species | 1 |
| Nomozaki-Akase |  | Nudibranchia | Doridacea(Cryptobranchia) |  | Chromodorididae |  | Doriprismatica | *Doriprismatica atromarginata*  (Cuvier,1804) | Tropical–subtropical–temperate species | 2 |
| Nomozaki-Akase |  | Nudibranchia | Doridacea(Porostomata) |  | Dendorodorididae |  | Dendrodoris | *Dendrodoris denisoni*  (Angas,1864) | Tropical–subtropical–temperate species | 1 |
| Nomozaki-Akase |  | Nudibranchia | Cladobranchia |  | Scyllaeidae |  | Scyllaea | *Scyllaea pelagica*(Linnaeus,1758) | Tropical–subtropical species | 1 |
| Nomozaki-Akase |  | Nudibranchia |  | Fionidea | Samlidae |  | Samla | *Samla takashigei*  Korshunova et al.,2017 | Tropical–subtropical species | 4 |
| Nomozaki-Akase | 2023-11-16 | Sacoglossa |  |  | Plakobranchidae |  | Thuridilla | *Thuridilla splendens*(Baba,1949) | Tropical–subtropical species | 1 |
| Nomozaki-Akase |  | Nudibranchia | Doridacea(Cryptobranchia) |  | Chromodorididae |  | Doriprismatica | *Doriprismatica atromarginata*  (Cuvier,1804) | Tropical–subtropical–temperate species | 1 |
| Nomozaki-Akase |  | Nudibranchia | Doridacea(Cryptobranchia) |  | Chromodorididae |  | Goniobranchus | *Goniobranchus tinctorius*  (Ruppell&Leuckart,1830) | Tropical–subtropical species | 1 |
| Nomozaki-Akase |  | Nudibranchia | Doridacea(Cryptobranchia) |  | Chromodorididae |  | Goniobranchus | *Goniobranchus orientails*  (Rudman,1983) | Tropical–subtropical species | 2 |
| Nomozaki-Akase |  | Nudibranchia | Doridacea(Porostomata) |  | Dendorodorididae |  | Dendrodoris | *Dendrodoris denisoni*  (Angas,1864) | Tropical–subtropical–temperate species | 1 |
| Nomozaki-Akase |  | Nudibranchia | Doridacea(Phanerobranchia) |  | Polyceridae |  | Polycerinae | *Polycern japonica* Baba,1949) | Tropical–subtropical–temperate species | 1 |
| Nomozaki-Akase |  | Nudibranchia | Cladobranchia |  | Tritoniidae |  | Tritoniopsis | *Tritoniopsis elegans*  (Audouin in Savigny,1826) | Tropical–subtropical species | 12 |
| Nomozaki-Akase |  | Nudibranchia |  | Fionidea | Samlidae |  | Samla | *Samla takashigei*  Korshunova et al.,2017 | Tropical–subtropical species | 4 |
| Nomozaki-Akase |  | Nudibranchia |  | Aeolidioidea | Facelinidae |  | Facelinella | *Facelinella anulifera* Baba,1949 | Tropical–subtropical species | 1 |
| Nomozaki-Akase | 2024-01-16 | Nudibranchia |  |  | Plakobranchidae |  | Elysia | *Elysia asbecki*  Wagele, Stemmer, Burghardt&Handeler,2010 | Tropical–subtropical species | 1 |
| Nomozaki-Akase |  | Sacoglossa |  |  | Plakobranchidae |  | Elysia | *Elysia atroviridis* Baba,1955 | Temperate species | 1 |
| Nomozaki-Akase |  | Sacoglossa |  |  | Plakobranchidae |  | Elysia | *Elysia* cf. *japonica* Eliot, 1913 | Temperate species | 2 |
| Nomozaki-Akase |  | Sacoglossa |  |  | Plakobranchidae |  | Elysia | *Elysia lobata* Gould,1852 | Tropical–subtropical species | 1 |
| Nomozaki-Akase |  | Sacoglossa |  |  | Elysiidae |  | Elysia | *Elysia marginata*(Pease,1871) | Tropical–subtropical–temperate species | 3 |
| Nomozaki-Akase |  | Sacoglossa |  |  | Plakobranchidae |  | Thuridilla | *Thuridilla splendens*(Baba,1949) | Tropical–subtropical species | 2 |
| Nomozaki-Akase |  | Sacoglossa |  |  | Plakobranchidae |  | Thuridilla | *Thuridilla vataae*(Risbec,1928) | Tropical–subtropical species | 2 |
| Nomozaki-Akase |  | Nudibranchia | Doridacea(Cryptobranchia) |  | Chromodorididae |  | Goniobranchus | *Goniobranchus tinctorius*  (Ruppell&Leuckart,1830) | Tropical–subtropical species | 2 |
| Nomozaki-Akase |  | Nudibranchia | Doridacea(Cryptobranchia) |  | Chromodorididae |  | Goniobranchus | *Goniobranchus orientails*  (Rudman,1983) | Tropical–subtropical species | 2 |
| Nomozaki-Akase |  | Nudibranchia | Doridacea(Cryptobranchia) |  | Chromodorididae |  | Hypselodoris | *Hypselodoris festiva*  (A.Adams,1861) | Temperate species | 2 |
| Nomozaki-Akase |  | Nudibranchia | Doridacea(Porostomata) |  | Dendorodorididae |  | Dendrodoris | *Dendrodoris denisoni*  (Angas,1864) | Tropical–subtropical–temperate species | 7 |
| Nomozaki-Akase |  | Nudibranchia | Doridacea(Phanerobranchia) |  | Polyceridae | Polycerinae | Polycera | *Polycera* sp.7 | Temperate species | 1 |
| Nomozaki-Akase |  | Nudibranchia | Doridacea(Phanerobranchia) |  | Goniodorididae |  | Goniodoris | *Goniodoris castanea* Alder&Hancock,1845 | Tropical–subtropical species | 1 |
| Nomozaki-Akase |  | Nudibranchia | Cladobranchia |  | Dotidae |  | Doto | *Doto japonica* Odhner,1936 | Temperate species | 1 |
| Nomozaki-Akase |  | Nudibranchia |  | Fionidea | Samlidae |  | Samla | *Samla takashigei*  Korshunova et al.,2017 | Tropical–subtropical species | 9 |
| Nomozaki-Akase |  | Nudibranchia |  | Aeolidioidea | Aeolidiidae |  | Bulbaeolidia | Bulbaeolidia alba(Risbec,1928) | Tropical–subtropical species | 1 |
| Tatsunoguchi | 2023-06-02 | Cephalaspidea |  |  | Aglajidae |  | Chelidonura | *Chelidonura hirundinina*  (Quoy&Gaimard,1833) | Tropical–subtropical–temperate species | 2 |
| Tatsunoguchi |  | Sacoglossa |  |  | Plakobranchidae |  | Elysia | *Elysia trisinuata* Baba,1949 | Tropical–subtropical species | 1 |
| Tatsunoguchi |  | Nudibranchia | Doridacea(Cryptobranchia) |  | Chromodorididae |  | Doriprismatica | *Doriprismatica atromarginata*  (Cuvier,1804) | Tropical–subtropical–temperate species | 1 |
| Tatsunoguchi |  | Nudibranchia | Doridacea(Cryptobranchia) |  | Chromodorididae |  | Goniobranchus | *Goniobranchus orientails*  (Rudman,1983) | Tropical–subtropical species | 3 |
| Tatsunoguchi |  | Nudibranchia | Doridacea(Cryptobranchia) |  | Chromodorididae |  | Goniobranchus | *Goniobranchus sinensis*  (Rudman,1985) | Tropical–subtropical species | 2 |
| Tatsunoguchi |  | Nudibranchia | Doridacea(Cryptobranchia) |  | Chromodorididae |  | Hypselodoris | *Hypselodoris sagamiensis*  (Baba,1949) | Tropical–subtropical species | 2 |
| Tatsunoguchi |  | Nudibranchia | Doridacea(Cryptobranchia) |  | Chromodorididae |  | Verconia | *Verconia nivalis*  (Baba,1937) | Temperate species | 1 |
| Tatsunoguchi |  | Nudibranchia | Doridacea(Cryptobranchia) |  | Discodorididae |  | Jorunna | *Jorunna parva*(Baba,1938) | Tropical–subtropical species | 2 |
| Tatsunoguchi | 2023-07-31 | Sacoglossa |  |  | Limapontiidae |  | Stiliger | *Stiliger ornatus* Ehrenberg,1828 | Tropical–subtropical species | 2 |
| Tatsunoguchi |  | Nudibranchia | Doridacea(Cryptobranchia) |  | Chromodorididae |  | Doriprismatica | *Doriprismatica atromarginata*  (Cuvier,1804) | Tropical–subtropical–temperate species | 9 |
| Tatsunoguchi |  | Nudibranchia | Doridacea(Cryptobranchia) |  | Chromodorididae |  | Goniobranchus | *Goniobranchus orientails*  (Rudman,1983) | Tropical–subtropical species | 1 |
| Tatsunoguchi |  | Nudibranchia | Doridacea(Cryptobranchia) |  | Chromodorididae |  | Goniobranchus | *Goniobranchus sinensis*  (Rudman,1985) | Tropical–subtropical species | 2 |
| Tatsunoguchi |  | Nudibranchia | Doridacea(Cryptobranchia) |  | Chromodorididae |  | Goniobranchus | *Goniobranchus fidelis*  (Kelaart,1858) | Tropical–subtropical–temperate species | 4 |
| Tatsunoguchi |  | Nudibranchia | Doridacea(Porostomata) |  | Phyllidiidae |  | Phyllidiella | *Phyllidiella pustulosa*(Cuvier,1804 | Tropical–subtropical–temperate species | 2 |
| Tatsunoguchi |  | Nudibranchia | Cladobranchia |  | Tritoniidae |  | Tritoniopsis | *Tritoniopsis elegans*  (Audouin in Savigny,1826) | Tropical–subtropical species | 3 |
| Tatsunoguchi | 2023-08-29 | Sacoglossa |  |  | Elysiidae |  | Elysia | *Elysia marginata*(Pease,1871) | Tropical–subtropical–temperate species | 2 |
| Tatsunoguchi |  | Nudibranchia | Doridacea(Cryptobranchia) |  | Chromodorididae |  | Doriprismatica | *Doriprismatica atromarginata*  (Cuvier,1804) | Tropical–subtropical–temperate species | 2 |
| Tatsunoguchi |  | Nudibranchia | Doridacea(Cryptobranchia) |  | Chromodorididae |  | Goniobranchus | *Goniobranchus fidelis*  (Kelaart,1858) | Tropical–subtropical–temperate species | 3 |
| Tatsunoguchi |  | Nudibranchia | Doridacea(Cryptobranchia) |  | Chromodorididae |  | Goniobranchus | *Goniobranchus orientails*  (Rudman,1983) | Tropical–subtropical species | 2 |
| Tatsunoguchi |  | Nudibranchia | Doridacea(Cryptobranchia) |  | Chromodorididae |  | Goniobranchus | *Goniobranchus sinensis*  (Rudman,1985) | Tropical–subtropical species | 1 |
| Tatsunoguchi |  | Nudibranchia | Doridacea(Cryptobranchia) |  | Chromodorididae |  | Cadlinella | *Cadlinella ornatissima*  (Risbec,1928) | Tropical–subtropical species | 1 |
| Tatsunoguchi |  | Nudibranchia | Doridacea(Cryptobranchia) |  | Chromodorididae |  | Verconia | *Verconia nivalis*(Baba,1937) | Temperate species | 1 |
| Tatsunoguchi |  | Nudibranchia | Doridacea(Cryptobranchia) |  | Chromodorididae |  | Goniobranchus | *Goniobranchus tinctorius*  (Ruppell&Leuckart,1830) | Tropical–subtropical species | 1 |
| Tatsunoguchi |  | Nudibranchia | Doridacea(Cryptobranchia) |  | Chromodorididae |  | Mexichromis | *Mexichromis multituberculata*  (Baba,1953) | Tropical–subtropical species | 1 |
| Tatsunoguchi |  | Nudibranchia | Cladobranchia |  | Tritoniidae |  | Tritoniopsis | *Tritoniopsis elegans*  (Audouin in Savigny,1826) | Tropical–subtropical species | 17 |
| Tatsunoguchi |  | Nudibranchia | Cladobranchia |  | Aminidae |  | Dematobranchus | *Dermatobranchus primus* Baba,1976 | Temperate species | 1 |
| Tatsunoguchi |  | Nudibranchia | Cladobranchia | Fionidea | Samlidae |  | Samla | *Samla takashigei*  Korshunova et al.,2017 | Tropical–subtropical species | 3 |
| Tatsunoguchi |  | Nudibranchia | Cladobranchia | Fionidea | Flabellinidae |  | Flabellina | *Flabellina* sp.3 | Tropical–subtropical species | 1 |
| Tatsunoguchi | 2023-09-19 | Sacoglossa |  |  | Elysiidae |  | Elysia | *Elysia marginata*(Pease,1871) | Tropical–subtropical–temperate species | 2 |
| Tatsunoguchi |  | Nudibranchia | Doridacea(Cryptobranchia) |  | Chromodorididae |  | Doriprismatica | *Doriprismatica atromarginata*  (Cuvier,1804) | Tropical–subtropical–temperate species | 9 |
| Tatsunoguchi |  | Nudibranchia | Doridacea(Cryptobranchia) |  | Chromodorididae |  | Goniobranchus | *Goniobranchus fidelis*  (Kelaart,1858) | Tropical–subtropical–temperate species | 1 |
| Tatsunoguchi |  | Nudibranchia | Doridacea(Cryptobranchia) |  | Chromodorididae |  | Goniobranchus | *Goniobranchus orientails*  (Rudman,1983) | Tropical–subtropical species | 1 |
| Tatsunoguchi |  | Nudibranchia | Doridacea(Cryptobranchia) |  | Chromodorididae |  | Verconia | *Verconia nivalis*(Baba,1937) | Temperate species | 1 |
| Tatsunoguchi |  | Nudibranchia | Doridacea(Cryptobranchia) |  | Chromodorididae |  | Cadlinella | *Cadlinella ornatissima*  (Risbec,1928) | Tropical–subtropical species | 1 |
| Tatsunoguchi |  | Nudibranchia | Cladobranchia |  | Tritoniidae |  | Tritoniopsis | *Tritoniopsis elegans*  (Audouin in Savigny,1826) | Tropical–subtropical species | 16 |
| Tatsunoguchi |  | Nudibranchia |  | Fionidea | Samlidae |  | Samla | *Samla takashigei*  Korshunova et al.,2017 | Tropical–subtropical species | 1 |
| Tatsunoguchi | 2023-10-23 | Sacoglossa |  |  | Limapontiidae |  | Stiliger | *Stiliger ornatus* Ehrenberg,1828 | Tropical–subtropical species | 1 |
| Tatsunoguchi |  | Nudibranchia |  |  | Chromodorididae |  | Goniobranchus | *Goniobranchus geometricus*  (Risbec,1928) | Tropical–subtropical species | 1 |
| Tatsunoguchi |  | Nudibranchia | Doridacea(Cryptobranchia) |  | Chromodorididae |  | Goniobranchus | *Goniobranchus fidelis*  (Kelaart,1858) | Tropical–subtropical–temperate species | 1 |
| Tatsunoguchi |  | Nudibranchia | Doridacea(Cryptobranchia) |  | Chromodorididae |  | Goniobranchus | *Goniobranchus sinensis*  (Rudman,1985) | Tropical–subtropical species | 2 |
| Tatsunoguchi |  | Nudibranchia | Doridacea(Cryptobranchia) |  | Chromodorididae |  | Goniobranchus | *Goniobranchus tinctorius*  (Ruppell&Leuckart,1830) | Tropical–subtropical species | 2 |
| Tatsunoguchi |  | Nudibranchia | Cladobranchia |  | Tritoniidae |  | Tritoniopsis | *Tritoniopsis elegans*  (Audouin in Savigny,1826) | Tropical–subtropical species | 2 |
| Tatsunoguchi |  | Nudibranchia | Cladobranchia |  | Aminidae |  | Dematobranchus | *Dermatobranchus primus* Baba,1976 | Temperate species | 4 |
| Tatsunoguchi | 2023-11-24 | Nudibranchia | Doridacea(Cryptobranchia) |  | Chromodorididae |  | Goniobranchus | *Goniobranchus tinctorius*  (Ruppell&Leuckart,1830) | Tropical–subtropical species | 1 |
| Tatsunoguchi | 2024-01-19 | Sacoglossa |  |  | Plakobranchidae |  | Elysia | *Elysia* cf. *japonica* Eliot, 1913 | Temperate species | 4 |
| Tatsunoguchi |  | Sacoglossa |  |  | Elysiidae |  | Elysia | *Elysia marginata*(Pease,1871) | Tropical–subtropical–temperate species | 1 |
| Tatsunoguchi |  | Sacoglossa |  |  | Plakobranchidae |  | Thuridilla | *Thuridilla splendens*(Baba,1949) | Tropical–subtropical species | 1 |
| Tatsunoguchi |  | Nudibranchia | Doridacea(Cryptobranchia) |  | Chromodorididae |  | Doriprismatica | *Doriprismatica atromarginata*  (Cuvier,1804) | Tropical–subtropical–temperate species | 2 |
| Tatsunoguchi |  | Nudibranchia | Doridacea(Cryptobranchia) |  | Chromodorididae |  | Goniobranchus | *Goniobranchus tinctorius*  (Ruppell&Leuckart,1830) | Tropical–subtropical species | 3 |
| Tatsunoguchi |  | Nudibranchia | Doridacea(Cryptobranchia) |  | Chromodorididae |  | Goniobranchus | *Goniobranchus fidelis*  (Kelaart,1858) | Tropical–subtropical–temperate species | 1 |
| Tatsunoguchi |  | Nudibranchia | Doridacea(Cryptobranchia) |  | Chromodorididae |  | Goniobranchus | *Goniobranchus orientails*  (Rudman,1983) | Tropical–subtropical species | 11 |
| Tatsunoguchi |  | Nudibranchia | Doridacea(Cryptobranchia) |  | Chromodorididae |  | Hypselodoris | *Hypselodoris sagamiensis*  (Baba,1949) | Tropical–subtropical species | 2 |
| Tatsunoguchi |  | Nudibranchia | Doridacea(Cryptobranchia) |  | Chromodorididae |  | Verconia | *Verconia nivalis*(Baba,1937) | Temperate species | 4 |
| Tatsunoguchi |  | Nudibranchia | Doridacea(Cryptobranchia) |  | Chromodorididae |  | Verconia | *Verconia purpurea*(Baba,1949) | Temperate species | 2 |
| Tatsunoguchi |  | Nudibranchia | Doridacea(Porostomata) |  | Dendorodorididae |  | Dendrodoris | *Dendrodoris denisoni*  (Angas,1864) | Tropical–subtropical–temperate species | 4 |
| Tatsunoguchi |  | Nudibranchia | Doridacea(Cryptobranchia) |  | Discodorididae |  | Jorunna | *Jorunna parva*(Baba,1938) | Tropical–subtropical species | 2 |
| Tatsunoguchi |  | Nudibranchia | Doridacea(Phanerobranchia) |  | Onchidoridoidae |  | Diaphodoris | *Diaphorodoris mitsuii*(Baba,1938) | Tropical–subtropical species | 4 |
| Tatsunoguchi |  | Nudibranchia | Doridacea(Phanerobranchia) |  | Goniodorididae |  | Okenia | *Okenia japonica* Baba,1949 | Temperate species | 3 |
| Tatsunoguchi |  | Nudibranchia | Cladobranchia |  | Aminidae |  | Dematobranchus | *Dermatobranchus primus* Baba,1976 | Temperate species | 7 |
| Tatsunoguchi |  | Nudibranchia | Cladobranchia |  | Tritoniidae |  | Tritoniopsis | *Tritoniopsis elegans*  (Audouin in Savigny,1826) | Tropical–subtropical species | 2 |
| Tatsunoguchi |  | Nudibranchia | Cladobranchia | Fionidea | Fionidae |  | Tenellia | *Tenellia* sp.44 | Tropical–subtropical species | 1 |
| Tatsunoguchi |  | Nudibranchia | Cladobranchia |  | Tritoniidae |  | Marionia | *Marionia* sp.1 | Temperate species | 1 |
| Tatsunoguchi |  | Nudibranchia |  | Aeolidioidea | Aeolidiidae |  | Bulbaeolidia | *Bulbaeolidia alba*(Risbec,1928) | Tropical–subtropical species | 1 |
| Tatsunoguchi |  | Nudibranchia |  | Aeolidioidea | Facelinidae |  | Caloria | *Caloria indica*(Bergh,1896) | Tropical–subtropical–temperate species | 1 |

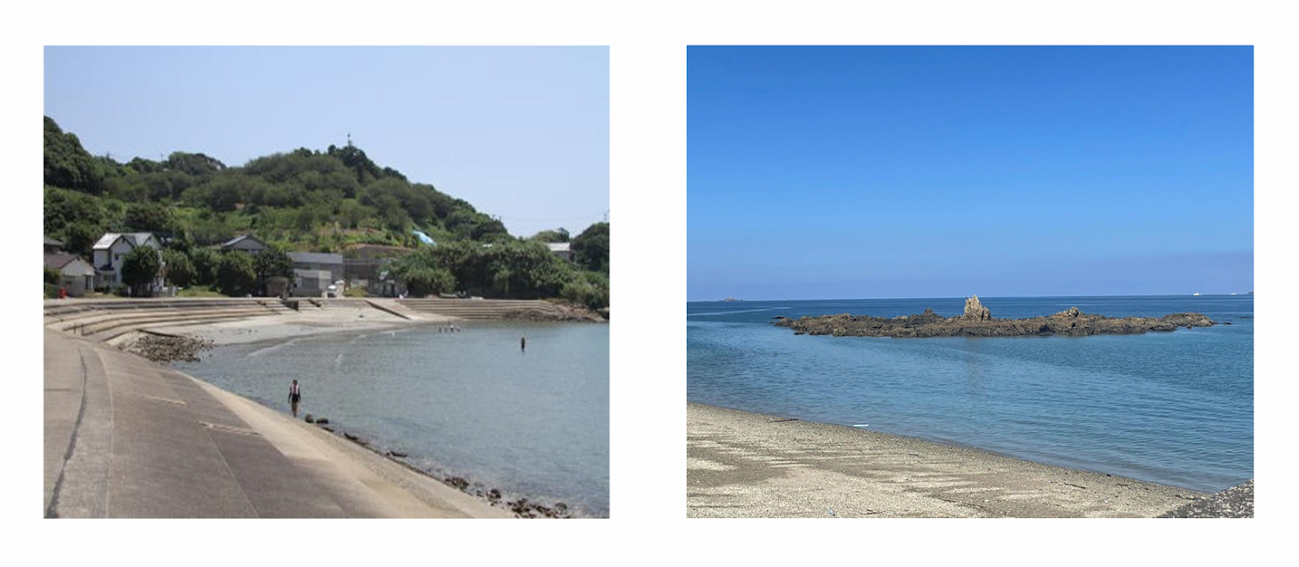

**Fig. S1** Survey sites used in this study and in Kawahara (unpublished). Left: Tatsunokuchi; Right: Nomozaki Akase. The present study area is shown in Fig. 1.

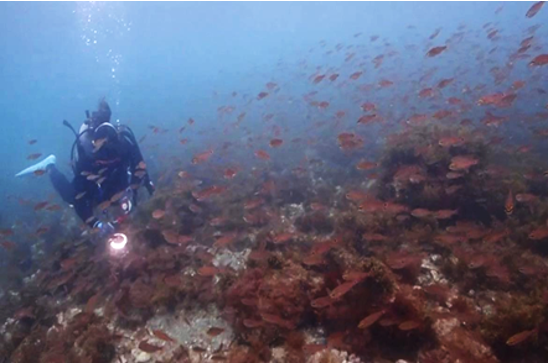

**Fig. S2** Photograph showing diving surveys conducted in this study.

**Fig. S3** Photographs illustrate the morphological diversity of nudibranchs recorded from coastal waters of northwestern Kyushu, Japan, during the 2023–2024 underwater surveys.

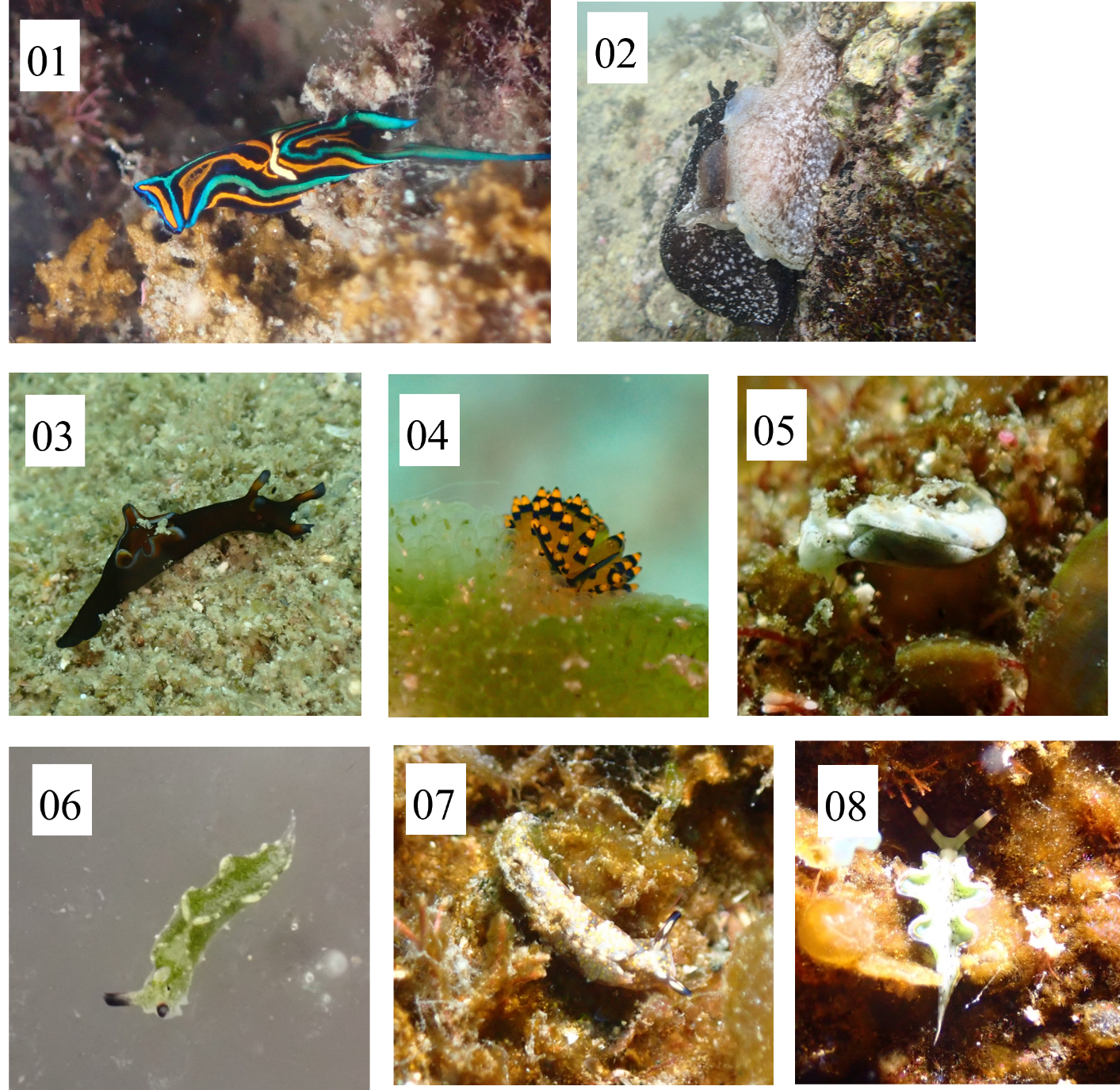
01.*Chelidonura hirundinina*(Quoy&Gaimard,1833)

02. *Aplysia kurodai*(Baba,1937)

03. *Aplysia japonica*(G.B.Sowerby Ⅱ,1869)

04. *Stiliger ornatus Ehrenberg*(Ehrenberg,1828)

05. *Elysia asbecki*(Wagele,Stemmer,Burghardt&Handeler,2010)

06. *Elysia atroviridis*（Baba,1955）

07. *Elysia* cf.j*aponica*(1913)

08. *Elysia lobata*(Gould,1852）

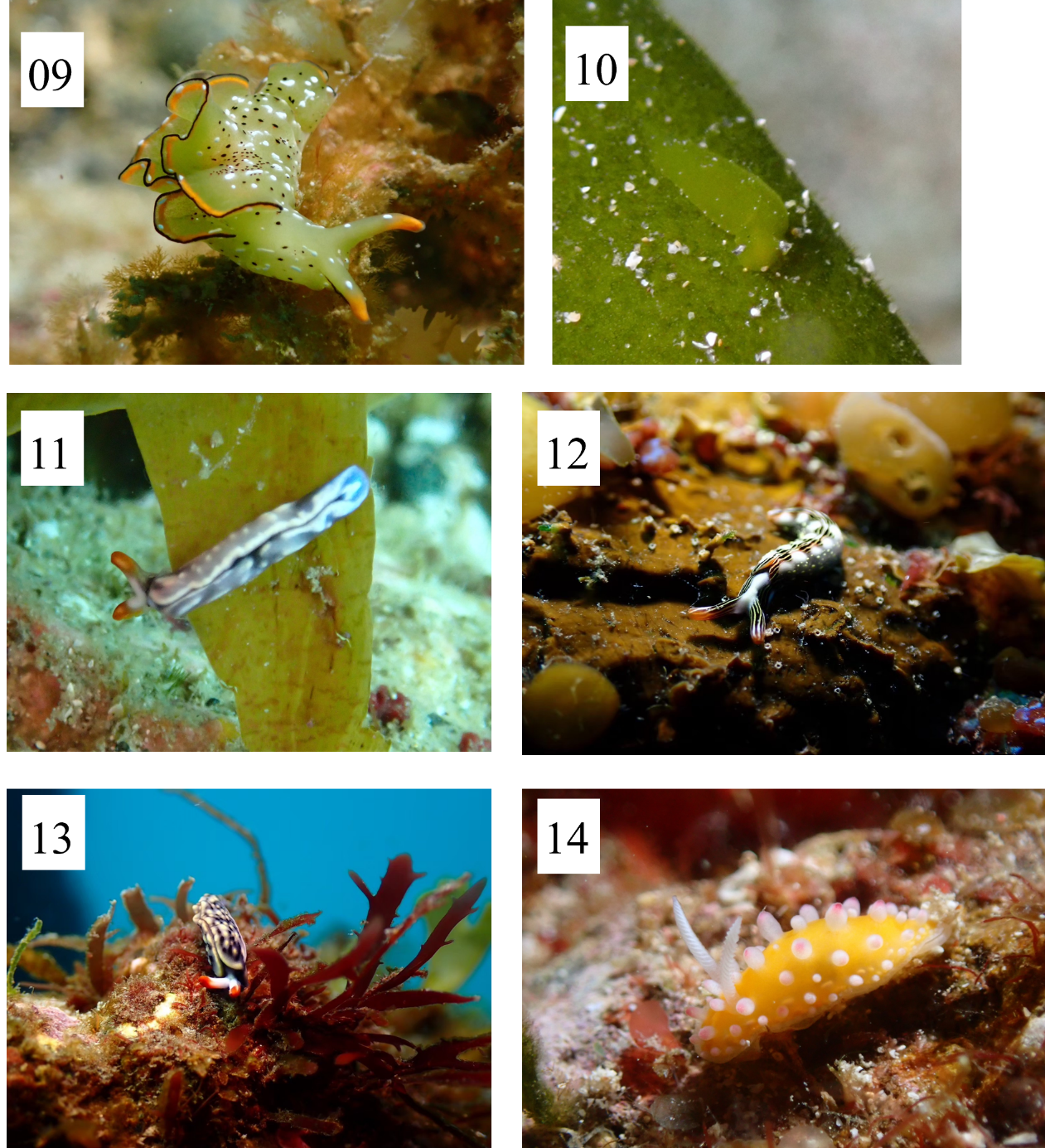

09. *Elysia marginata*(Pease,1871)

10. *Elysia trisinuata*(Baba,1949)

11. *Thuridilla albopustulosa*(Gosliner,1995)

12. *Thuridilla splendens*(Baba,1949)

13. *Thuridilla vataae*（Risbec,1928)

14. *Cadlinella ornatissima*(Risbec,1928)

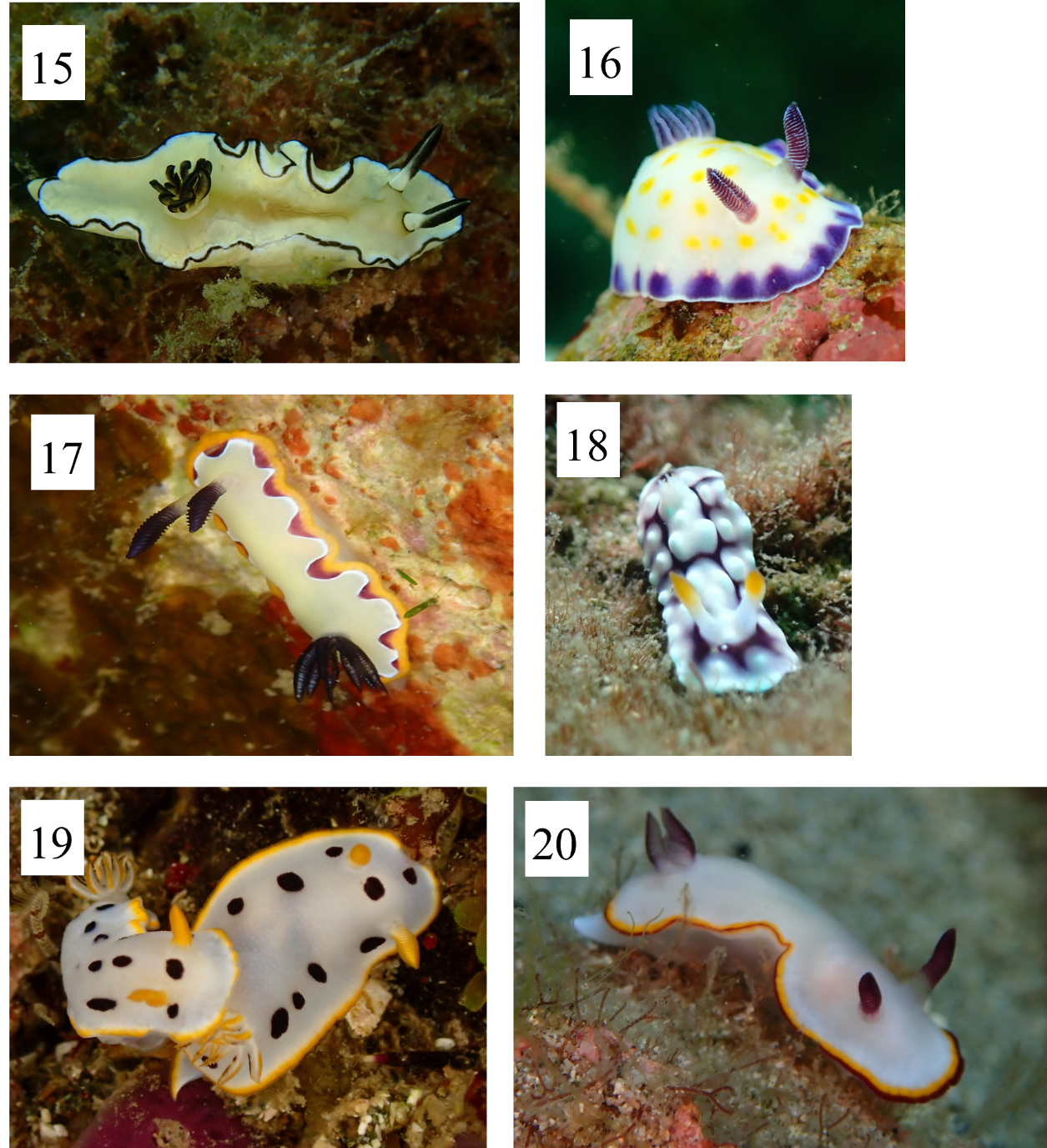

15. *Doriprismatica atromarginata*(Cuvier,1804)

16. *Goniobranchus aureopureus*(Collingwood,1881)

17. *Goniobranchus fidelis*(Kelaart,1858)

18. *Goniobranchus geometricus*

19. *Goniobranchus orientails*(Rudman,1983)

20. *Goniobranchus sinensis*(Rudman,1985)

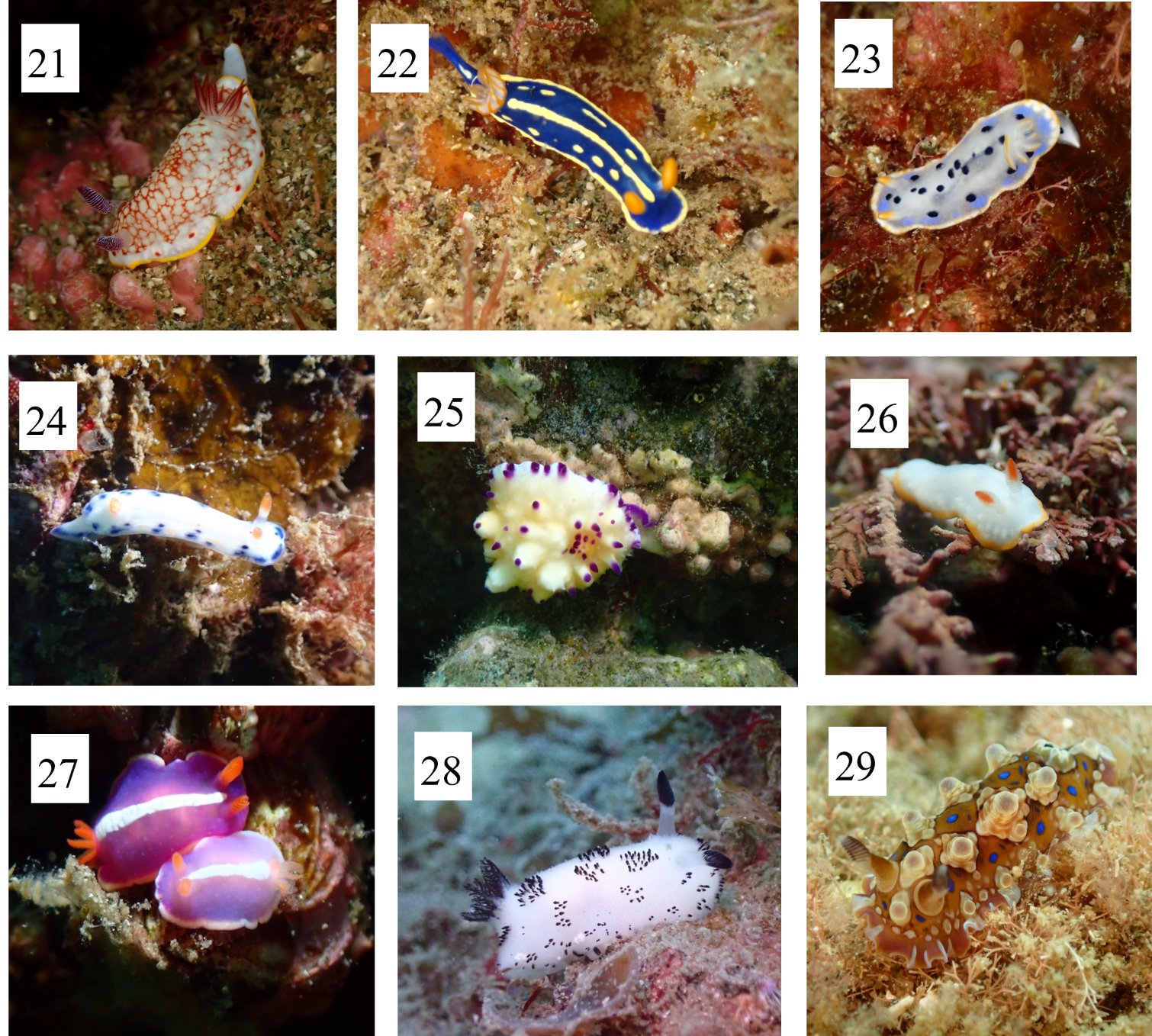

21. *Goniobranchus tinctorius*(Ruppell&Leuckart,1830)

22. *Hypselodoris festiva*(A.Adams,1861)

23. *Hypselodoris placida*(Baba,1949)

24. *Hypselodoris sagamiensis*(Baba,1949)

25. *Mexichromis multituberculata*(Baba,1953)

26. *Verconia nivalis*(Baba,1937)

27. *Verconia purpurea*(Baba,1949)

28. *Jorunna parva*(Baba,1938)

29. *Dendrodoris denisoni*(Angas,1864)

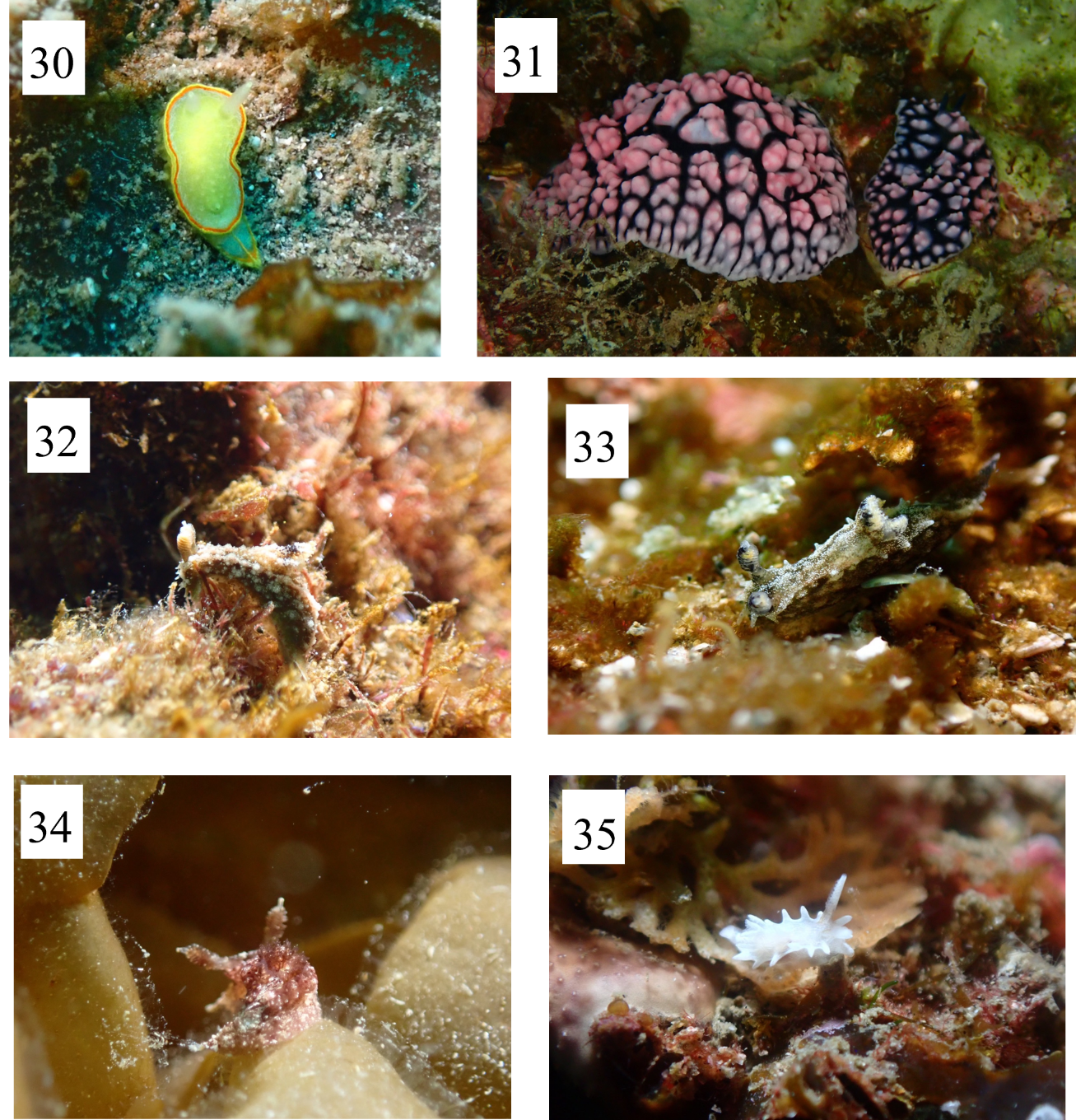

30. *Diaphorodoris mitsuii*

31. *Phyllidiella pustulosa*(Cuvier,1804)

32. *Polycera japonica*(Baba,1949)

33. *Polycera* st.7

34. *Pelagella castanea*(Alder&Hancock,1845)

35. *Okenia japonica*(Baba,1949)

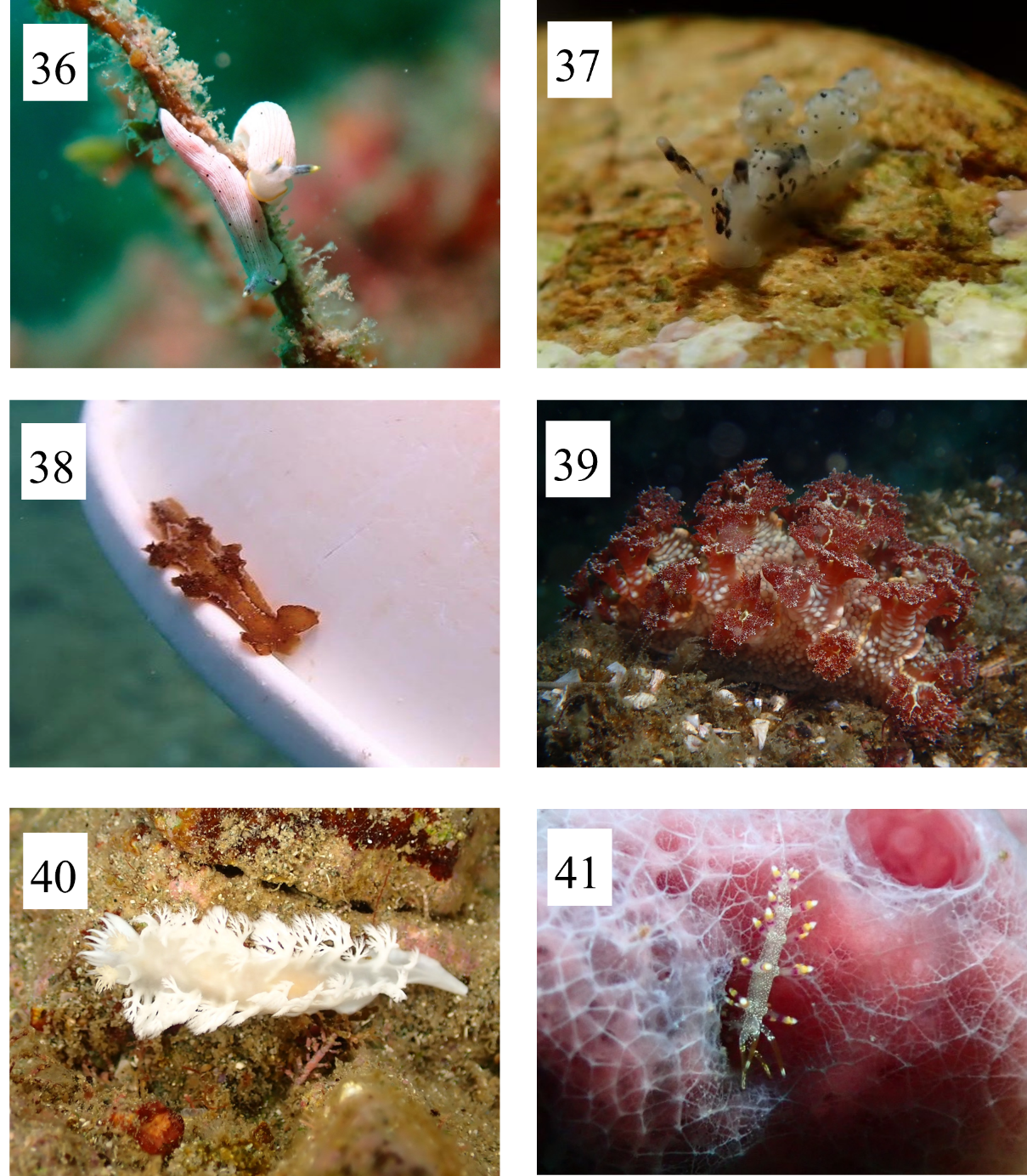

36. *Dermatobranchus primus* (Baba,1976)

37. *Doto japonica* (Odhner,1936)

38. *Scyllaea pelagica*(Linnaeus,1758)

39. *Marionia* sp.1

40. *Tritoniopsis elegans*(Audouin in Savigny,1826)

41. *Flabellia* sp.3

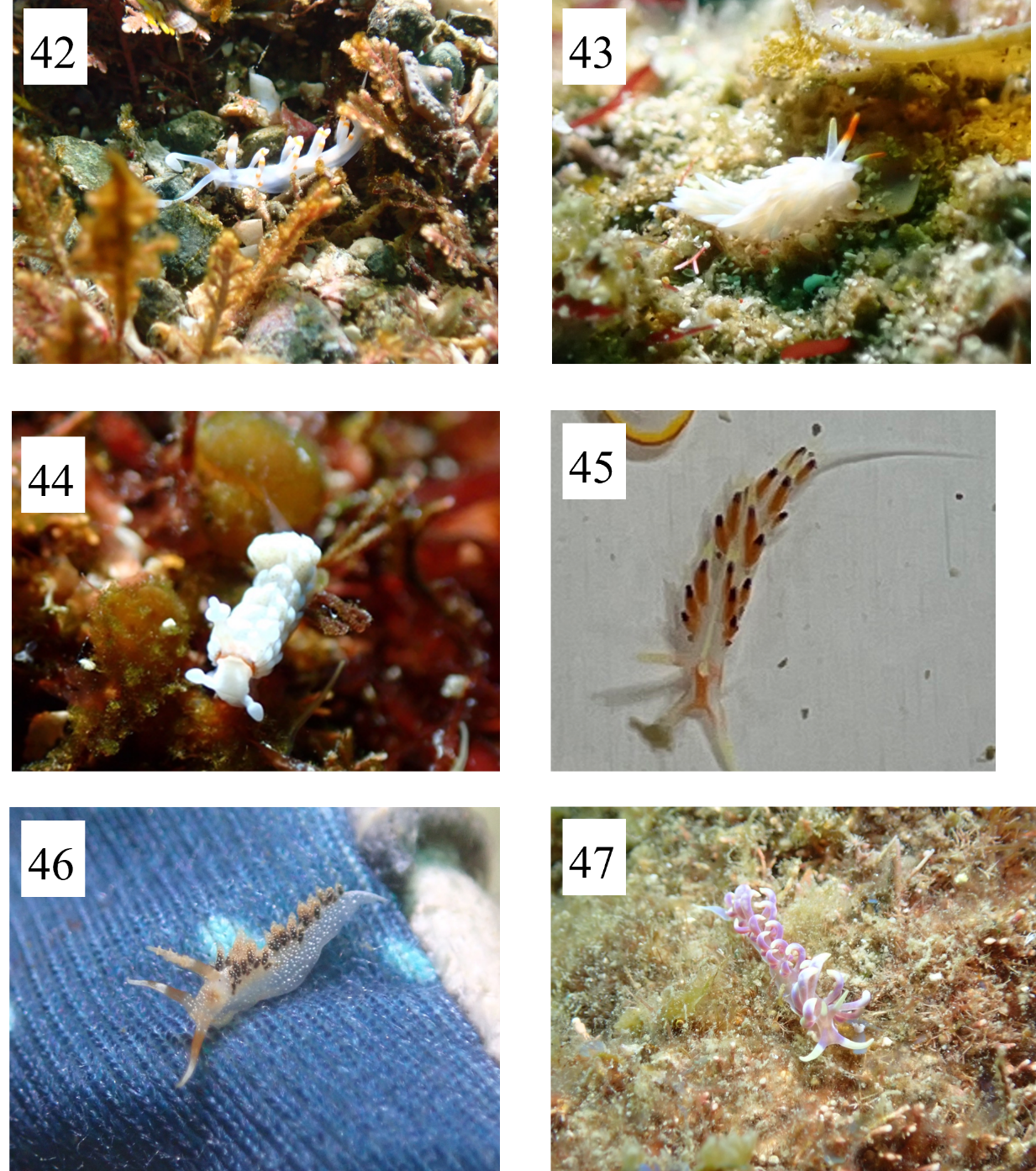

42. *Samla takashigei*(Korshunova et al.2017)

43. *Tenellia* sp.44

44. *Bulbaeolidia alba* (Risbec,1928)

45. *Caloria indica*(Bergh,1896)

46. *Facelinella anulifera*(Baba,1949)

47. *Phyllodesmium*(Rudman,1991)
